# Risk preferences causally rely on parietal magnitude representations: Evidence from combined TMS-fMRI

**DOI:** 10.1101/2025.01.13.632678

**Authors:** Gilles de Hollander, Marius Moisa, Christian C. Ruff

## Abstract

Risk preferences have traditionally been considered as stable traits that reflect subjective-valuation processes in prefrontal areas. More recently, however, it has been suggested that risk preferences may also be shaped by how choice problems are perceived and processed in perceptual brain regions. Specifically, the acuity of the parietal approximate number system (ANS), which encodes payoff magnitudes for different choice options, has been shown to correlate with both risk preferences and choice consistency. However, this correlational relationship leaves open the question whether parietal magnitude representations in fact causally underlie choice. Here, we provide direct evidence for such a key causal role of parietal magnitude representations in economic choice, using continuous theta-burst transcranial magnetic stimulation (cTBS), combined with functional MRI (fMRI) and numerical population receptive field (nPRF) modeling.

Our stimulation protocol targeted numerosity-tuned regions in the right parietal cortex, identified through nPRF modeling of individual fMRI data (n=35; within-subject design). The stimulation successfully perturbed neural processing, as evidenced by decreased amplitude of numerical magnitude-tuned responses and less accurate multivariate decoding of presented magnitudes from unseen data that were not used for model fitting. In line with a perceptual account of risky choice, the reduction in neural information capacity was also reflected in noisier behavioral responses. Moreover, a computational cognitive model fitted to choice behavior revealed that perturbing the ANS specifically increased the noisiness of small-magnitude representations. This perturbation made small magnitudes to be perceived as larger than they actually are, leading to more risk-seeking behavior. Finally, individual estimates of the cTBS effect on cognitive noise correlated with the corresponding decrease in amplitude of numerical magnitude-tuned fMRI responses, further solidifying the role of the parietal ANS in economic choice. In conclusion, our study demonstrates that the precision of parietal magnitude representations causally influences economic decision-making, with noisier encoding promoting biased risk-taking as formalized in recent perceptual models of risky choice.

## Introduction

Risk attitudes reflect the extent to which decision-makers are willing to accept higher variance in returns (i.e., higher risk) in exchange for potentially greater rewards. For instance, stocks typically yield higher average returns than bonds but also carry a greater likelihood of losing value. Some investors prefer the stability of bonds, while others favor the higher potential gains of stocks. Similarly, some may prefer comprehensive car insurance when renting a vehicle, whereas others are comfortable opting for higher deductibles at a lower price. Many economic theorists view such risk attitudes as an individual trait: just as some people particularly dislike electronic music, others may particularly dislike taking risks (Stigler and Becker, 1977).

Individual risk attitudes are generally inferred from preferences, as elicited through tasks requiring choices between, for example, uncertain gambles and smaller but guaranteed payoffs (Mata et al., 2018; Pedroni et al., 2017). These behavioral preferences are then interpreted as proxies for underlying risk attitudes, revealing how individuals generally balance potential rewards and uncertainty. However, growing evidence suggests that risk preferences are not as static as one would expect from a stable trait. Experimenters can profoundly manipulate risk preferences by emphasizing different information (Bordalo et al., 2012; de Hollander et al., 2024; Molter & Mohr, 2021; Thomas et al., 2019), directing attention to specific aspects of a decision (Busemeyer & Townsend, 1993; Fiedler & Glöckner, 2012), or altering the range of payoffs participants have previously encountered (Stewart et al., 2006; Walasek & Stewart, 2015).

One explanation for the observed instability of behavioral risk preferences is that they may be influenced by our moment-to-moment perception and representation of choice problems, which is inherently noisy and highly sensitive to context and expectations (Khaw et al., 2020; Woodford, 2020). Just as we can misjudge distances or time intervals in perceptual tasks by adapting our perception towards what we have recently observed (Petzschner et al., 2015), overestimating small and underestimating large values, we might similarly misjudge the size of monetary payoffs in economic decision-making. For example, people might perceive relatively larger and riskier payoff magnitudes as smaller than they actually are, making their behavior risk-averse. This regressive bias is far from arbitrary but reflects an optimal Bayesian strategy: When information is uncertain or noisy, our brains rely more on prior beliefs (“which potential payoffs are plausible in the first place”) to reduce errors. While this can amplify systematic biases, it ultimately helps us make more accurate decisions over time and has consistently been observed in a wide array of perceptual domains (Petzschner et al., 2015).

In line with a potential role for variable perception during risky choice, the perception and neural representation of different types of magnitudes—such as sizes, payoffs, and probabilities, which are key factors in economic decision-making—have been linked to a specific cortical region in and around the intraparietal sulcus (IPS). This region exhibits tuned responses to numerosity, as well as other abstract magnitudes (Bueti & Walsh, 2009; Cai et al., 2023; Foucault et al., 2024; Harvey, 2016; Harvey et al., 2013, 2015, 2020) and has been identified as the neural locus of the approximate number system (ANS), a cognitive system involved in quick judgments of numerical magnitudes (ANS; Dehaene, 2011; Dehaene et al., 1998). Could the fidelity and distortions of these parietal neural representations of the key numerical determinants of a choice problem be driving some of the economic choices we make?

Multiple lines of recent evidence support such a perceptual account of risk preferences. First, individual differences in risk preferences are correlated with performance on perceptual tasks involving numerical magnitudes (Barretto-García et al., 2023; Peters et al., 2005). For example, individuals with lower acuity in judging numerosities during the perceptual task tend to be less consistent and more risk-averse when making decisions about risky prospects. This finding suggests that a single trait may underpin both the fidelity of rapid numerical judgments and the degree of behavioral bias exhibited in economic decision-making, aligning with a Bayesian perceptual framework for risky choices where increased perceptual noise leads to more biased estimates (overestimating the smaller payoffs that come with safe options and underestimating those of riskier ones). This bias results in economic choices that deviate from risk-neutrality. Critically, this bias seems to relate to activity in the parietal ANS: The accuracy of a multivariate decoder trained to predict numerical magnitudes from fMRI activity patterns in the right IPS has been shown to correlate with the variability and bias of both perceptual and economic choice behavior, pointing out the potentially critical role of the acuity of the parietal ANS in shaping biases that determine risk preferences (Barretto-García et al., 2023). Indeed, neurally, participants with a higher gray matter volume in the right IPS behave more like an ‘unbiased’ risk-neutral decision-maker as well (Gilaie-Dotan et al., 2014).

A second converging line of evidence revealed that temporal fluctuations in the acuity of neural magnitude representations in the right parietal ANS of *a single individual* predict their economic choices across decisions as well (de Hollander, Grueschow, et al., 2024). Decisions during which fMRI activity patterns in numerically-tuned parietal cortex were less consistent were associated with greater variability in participants’ choices. Computational modeling of behavior revealed that participants seemingly relied more heavily on prior beliefs when noise increased, as reflected in responses that deviated more from risk-neutrality when neural representations appeared noisier.

These earlier studies suggest a key role for magnitude representations in the parietal ANS and showcase the relevance of its properties for understanding risk preferences. However, studies on this topic have so far been exclusively correlational, raising the critical question of whether magnitude signals in the parietal ANS play a causal role in economic decision-making, or whether they merely reflect neurocognitive representations associated with choice. Without causal evidence, it remains possible that a factor outside the parietal cortex drives both behavior and the precision of parietal magnitude representations. For example, inter- and intraindividual differences in choice behavior might also be directly driven by attentional, arousal, or emotional traits/states (Jahedi et al., 2017; Olschewski et al., 2018; Stanton et al., 2014), so that the increased fidelity of neurocognitive representations in parietal cortex may be an epiphenomenon rather than than a neural origin of behavior (Pfeffer et al., 2022; Whitmarsh et al., 2022).

Notably, one study has already shown that perturbation of the left IPS using brain stimulation can modulate risky choices under some circumstances (Coutlee et al., 2016). Specifically, when choosing between a certain amount and a risky 50-50 gamble, participants were less likely to choose the risky option after IPS perturbation. However, this study did not look into the neural effects of the stimulation, nor did it investigate whether the reduced number of risky choices was the result of decreased choice consistency, a shift in average risk preferences, or a combination of both. Importantly, general alterations in decision-making noisiness (without any link to preferences) can manifest as either increased or decreased proportions of risky choices (Olschewski et al., 2022). Thus, a precise, mechanistic characterization of the causal role of the parietal ANS in risky choice is lacking.

Here, we seek to advance our understanding of the neural origins of risk preferences, by temporally perturbing the parietal approximate number system (ANS) in human participants during a choice task involving risky prospects with brain stimulation. Critically, we measured how this perturbation influenced neural representations of payoff magnitudes using fMRI as well. Furthermore, we employed a computational behavioral modeling framework to separately quantify the effects of the perturbation on both choice consistency and average risk preferences.

We used continuous theta-burst transcranial magnetic stimulation (cTBS; an offline transcranial magnetic stimulation (TMS) protocol applied before task performance Huang et al., 2005). Previewing our results, stimulation reduced the fidelity of parietal tuning to the presented payoff magnitudes. Moreover, we found reduced choice consistency and increased risk-seeking behavior, but only for trials where participants were presented with smaller, safe payoffs first. Individuals showing larger reductions in choice consistency due to cTBS also exhibited greater shifts in risk preferences, aligning with predictions from a Bayesian perceptual framework of decision-making in which increased noise leads to increased weighting of prior beliefs and stronger perceptual biases. Computational modeling of the cognitive processes underlying these effects confirmed that the shifts in preference and response consistency can be attributed to increased noise in processing smaller but not larger magnitudes. Finally, the model-based estimate of increased noise in magnitude processing correlated with reduced amplitudes of neurophysiological payoff representations (nPRFs) after stimulation, solidifying the tight link between the parietal ANS and choice behavior.

Together, these results provide critical insights into how neurocognitive representations of payoff magnitudes, localized in the parietal ANS, shape economic decisions and why people sometimes exhibit more or less risk-averse behavior. By identifying the role of representational parietal noise in influencing choice consistency and risk preferences, our study also sheds more light on the neuro-cognitive mechanisms underlying variability in decision-making under uncertainty over and above the presumed role for the ‘prefrontal’ valuation network (Bartra et al., 2013; Levy & Glimcher, 2012).

## Results

### Experimental approach

Our experiment consisted of three sessions, during which participants repeatedly performed a decision-making task involving risky prospects. The first session aimed to identify participants with the most clearly defined magnitude-tuned regions in the right intraparietal area. This clear definition was crucial because we aimed to specifically target numerically-tuned subregions of the parietal cortex rather than a location based on macroanatomy (via, e.g., MNI space). Thus, using fMRI, we assessed which subregions of the parietal cortex exhibited sufficiently reliable tuning to specific stimulus numerosities within the relatively short scanning time permitted by our TMS design, and confirmed that these subregions were accessible for subsequent stimulation (i.e., didn’t lie deep in the sulcus).

Participants who met pre-defined selection criteria (see Fig.1 and Methods for details) proceeded to two additional sessions. In these sessions, participants again performed the decision-making task, but with either active stimulation targeting the most reliably numerically tuned regions of the right parietal number area or sham stimulation applied to the vertex.

**Figure 1:**
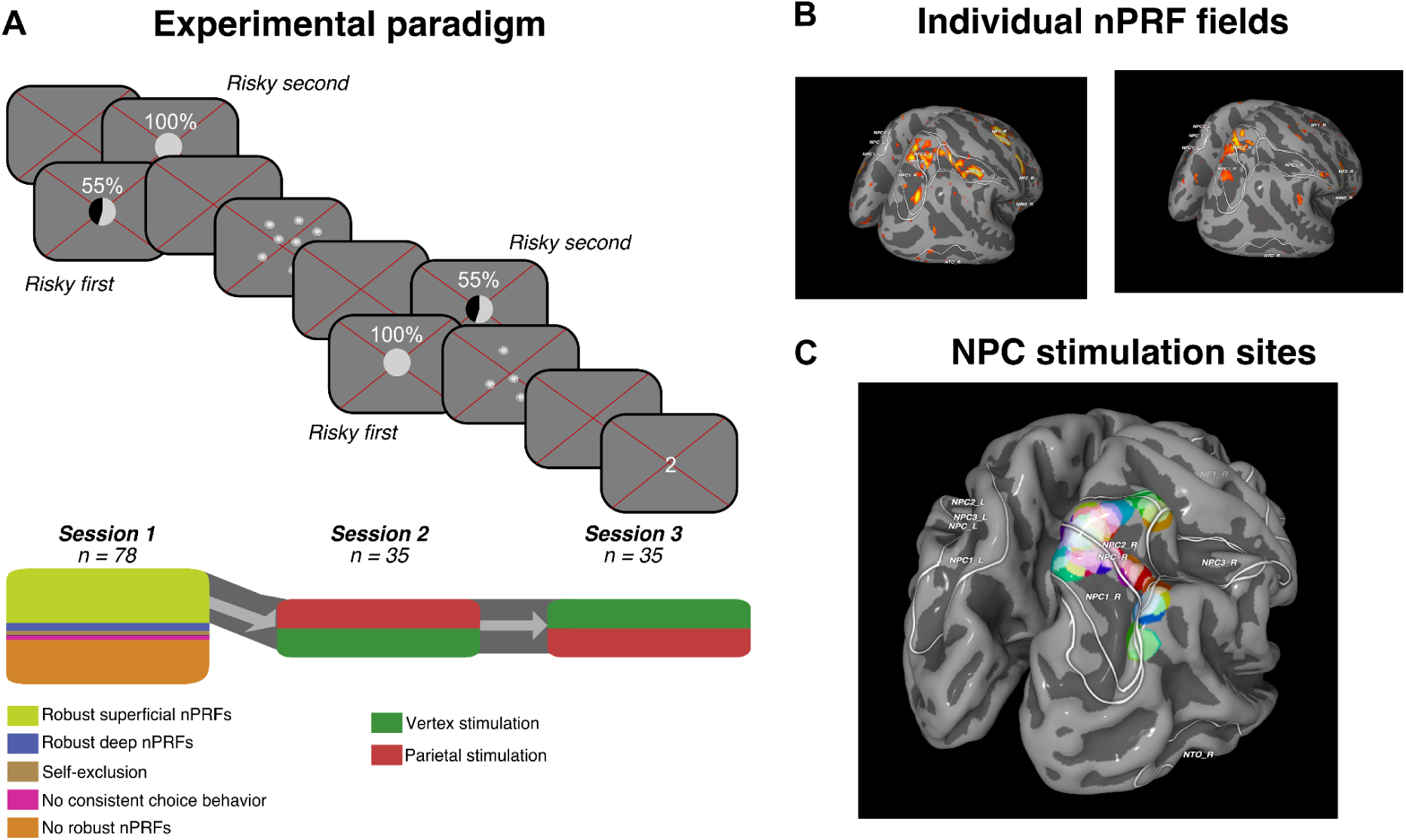
A) Experimental paradigm. Top panel: Participants were sequentially presented with two choice options: A larger payoff with a 55% payout probability or a smaller,payoff that was 100% certain. Presentation order was counterbalanced across trials (Same number of *Risky first* or *Risky second*) to control for previously established order effects. Participants could respond as soon as the second stimulus was presented and got feedback about which option they had chosen (but not the outcome of the gamble). Bottom panel: The first session was used to select participants for reliable numerically-tuned activation patterns in parietal cortex, as well as non-random behavior and TMS feasibility. **B) Individual nPRF maps**. Two representative examples of the explained variance maps. Note how robust numerosity tuning is mostly confined to areas in and around the lateral intraprietal area. **C) NPC stimulation sites.** Based on the estimated nPRF R2 maps, we selected a cTBS target on the rostral bank of the intraparietal sulcus. This figure shows all 35 targets as individual colored circles in *fsaverage*-space. Note that there are substantial interindividual differences in peak R2 clusters and therefores stimulation sites. This is in line with earlier work focusing on numerical (Harvey et al., 2013) and visual receptive fields in intraparietal cortex (Wang et al., 2015) that showed very substantial interindividual variability in the macro-anatomical location of functionally-defined regions.

The decision-making task, performed during both sessions while undergoing fMRI, consisted of 120 choices between a sure option with a fixed payout and a risky option with a 55% probability of a substantially larger payout. Options were presented sequentially (see Fig. 1A). The presentation order (i.e., risky option first or safe option first) was counterbalanced across trials since earlier work has revealed a strong order effect in this experimental paradigm related to working memory effects (de Hollander, Grueschow, et al., 2024). Participants were financially incentivized: One choice was randomly chosen and its outcome determined and paid out for each session.

Seventy-eight (78) participants (47 male, average age 23.9, ranged 18-35) joined the first experimental session of the experiment, of which thirty-five (35) subjects (23 male, average age 23.1, ranged 18-30) were selected for the two follow-up sessions involving TMS (see Methods for more details). Stimulation was conducted on the scanner bed, immediately before participants entered the scanner bore to perform the experimental paradigm. On average, 3 minutes and 13 seconds (std. 19 seconds) elapsed between the end of the stimulation protocol and the start of the first trial (see methods for more details).

Numerically tuned areas were identified by fitting a numerical receptive field (nPRF) model to single-trial fMRI responses to the first payoff magnitude. This early signal, preceding any decision-related activity from the second stimulus, isolates perception-related activity free from decision-making or response time effects (Barretto-García et al., 2023; de Hollander, Grueschow, et al., 2024; Harvey et al., 2013). The nPRF model characterizes voxels with reliable tuning to specific numerosities, showing peak activity for preferred values and decreasing activity for neighboring numerosities. Such tuned responses have been shown to predict individual differences in decision-making involving numerical magnitudes (Barretto-García et al., 2023; de Hollander, Grueschow, et al., 2024; Eger et al., 2009; Harvey et al., 2013; Lasne et al., 2019).

### Decreased amplitudes of numerically-tuned parietal cortex after cTBS

First, we probed the influence of cTBS on the noisiness of the *neural* representations of payoff magnitudes in the parietal ANS, by measuring its effect on observed nPRF parameters. The precise ROI we used was an individualized numerical parietal cortex mask that was defined as all voxels that (a) were within 2 centimeters of the center of the nPRF cluster from session one that was used to target the TMS coil (Fig. 1C), (b) fell within a predefined anatomical mask of the right numerical parietal cortex (NPC1 and NPC2 mask in Barretto-García et al., 2023), and (c) showed a cross-validated out-of-sample R^2^ higher than 0 in at least one of the two experimental sessions (i.e., session 2 or 3).

We hypothesized that cTBS should reduce neural responsivity, which would correspond to the nPRF model’s amplitude parameter. Indeed, we found significantly lower nPRF amplitudes after parietal stimulation (from 1.30 to 1.04 percent signal change, t(34) = 1.99, p=0.027, one-sided; Fig. 2B) compared to vertex stimulation. There was no difference in the average preferred numerosity (17.8 to 14.8; t(34) = 1.00, p=0.32), nor the dispersion of the nPRFs (from 0.9 to 0.77 sd in log space; t(34) = 1.27, p=0.21), highlighting a specific effect of TMS on the amplitude of the nPRF but not its numerosity tuning. The resulting activity patterns were also noisier, as reflected in the significantly reduced explained variance of the selected voxels (from 6.8% to 4.9%; t(34) = 2.06, p=0.023, one-sided). Furthermore, the total proportion of voxels that showed an out-of-sample-explained variance of more than 0 also decreased after cTBS on parietal cortex (from 11.1% to 7.5%, t(34) = 1.99, p=0.027, one-sided).

**Figure 2.**
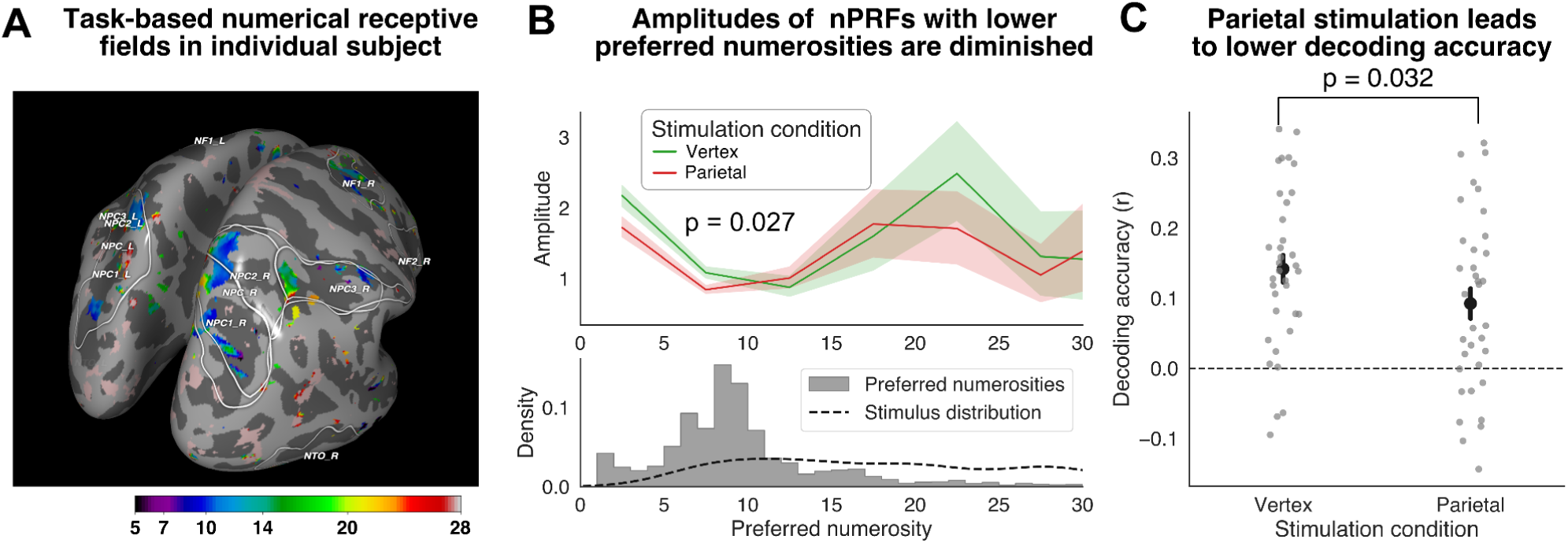
A) Preferred numerosities of numerically tuned cortical areas in a representative participant. reveal clear non-linear tuning around the intraparietal and postcentral sulcus, consistent with prior findings (Barretto-García et al., 2023; de Hollander, Grueschow, et al., 2024). **B) Responses amplitudes for vertex and parietal stimulation in numerosity-tuned parietal areas.** The amplitude of numerosity-tuned populations in parietal cortex was reduced after cTBS. Also, note how the majority of tuned populations had a relatively low preferred numerosity. Shaded areas correspond to the standard error of the mean over participants. **C) Decoding accuracy**: When the nPRF model was inverted to decode presented numerosities, accuracy decreased following cTBS in the parietal cortex (dots are individual differences, error bars standard error of the mean).

In line with our earlier work (Barretto-García et al., 2023; de Hollander, Grueschow, et al., 2024), we also inverted the nPRF model for individual participants and decoded the presented payoff magnitudes on a trial-by-trial basis from out-of-sample fMRI activity patterns. We hypothesized that the attenuation of parietal numerosity-tuned neural populations would reduce the fidelity of the neurocognitive representation, thereby reducing our ability to decode the objective numerosities from neural data. Indeed, the correlation between the decoded potential payoff and the actual payoff was lower after parietal ANS stimulation than vertex stimulation (mean r=0.142 versus r=0.092, F(1,34) = 4.99, p=0.032; Fig. 2C), in a similar manner for trials where risky or safe options were presented first (interaction of cTBS and presentation order: F(1, 34)=0.86, p=0.360).

In sum, the neural analysis confirmed that our cTBS protocol was successful in perturbing neural processing in numerically-tuned parietal cortex: The average amplitude in response to the stimuli was diminished, thereby also reducing the signal-to-noise ratio of these representations, but without changing their average tuning or specificity.

### Stimulation of parietal ANS decreases choice consistency and shifts average risk preference

A key behavioral hypothesis that follows from our perceptual framework is that the perturbation of neurocognitive representations of the payoff magnitudes should lead to less consistent choice behavior, because when neurocognitive representations become more variable, so should the resulting choice behavior.

To test this hypothesis, we fitted a psychophysical (probit) model using hierarchical Bayesian estimation (Barretto-García et al., 2023; Khaw et al., 2020; Olschewski et al., 2018, 2022; Olschewski & Rieskamp, 2021). This model predicts the mean proportion of risky choices for a given choice problem as a function of the log ratio between the payoff of the risky and safe option (accounting for Weber’s law; Barretto-García et al., 2023; de Hollander, Grueschow, et al., 2024; Khaw et al., 2020). The model assumes that the larger this ratio is, the more likely a participant is to choose the risky option. The strength of this relationship is a measure of the consistency of choices (and a direct analogue of psychophysical sensitivity in perceptual tasks; Green & Swets, 1966). The model also has an intercept parameter that (when scaled by the slope parameter) determines where the indifferent point of a participant lies – the ratio between risky and safe payoffs where the participant chooses the risky option 50% of the trials and which is our measure of average risk preference (see also Khaw et al., 2020). Note that we refrained from analyzing raw choice proportions since they cannot be interpreted cleanly in isolation (Olschewski & Rieskamp, 2021).

In our psychophysical model, we also included the order in which options were presented in our model (i.e., risky or safe option presented first). We did this because earlier work using the same experimental paradigm (de Hollander, Grueschow, et al., 2024) has revealed a profound effect of presentation order on average risk preference, reflecting that first-presented options in working memory are noisier and therefore perceived with more bias. Here, we replicated this order effect as well (risk-neutral probability^1^ was 51.0% [44.4, 57.0] when the risky option was presented first and 55.2% [47.5, 63.0] when the risky option came second; p_Bayesian_ = 0.001).

Our psychophysical analysis (Fig. 3) revealed that choice consistency was indeed significantly reduced after cTBS, but only for trials where the safe option was presented first (slope from 2.56 [2.19, 2.97] to 2.19 [1.83, 2.56]; p_Bayesian_=0.013), not when the risky option was presented first (slope from 2.44 [2.05, 2.85] to 2.41 [2.19, 2.97], p_Bayesian_=0.446).

**Figure 3.**
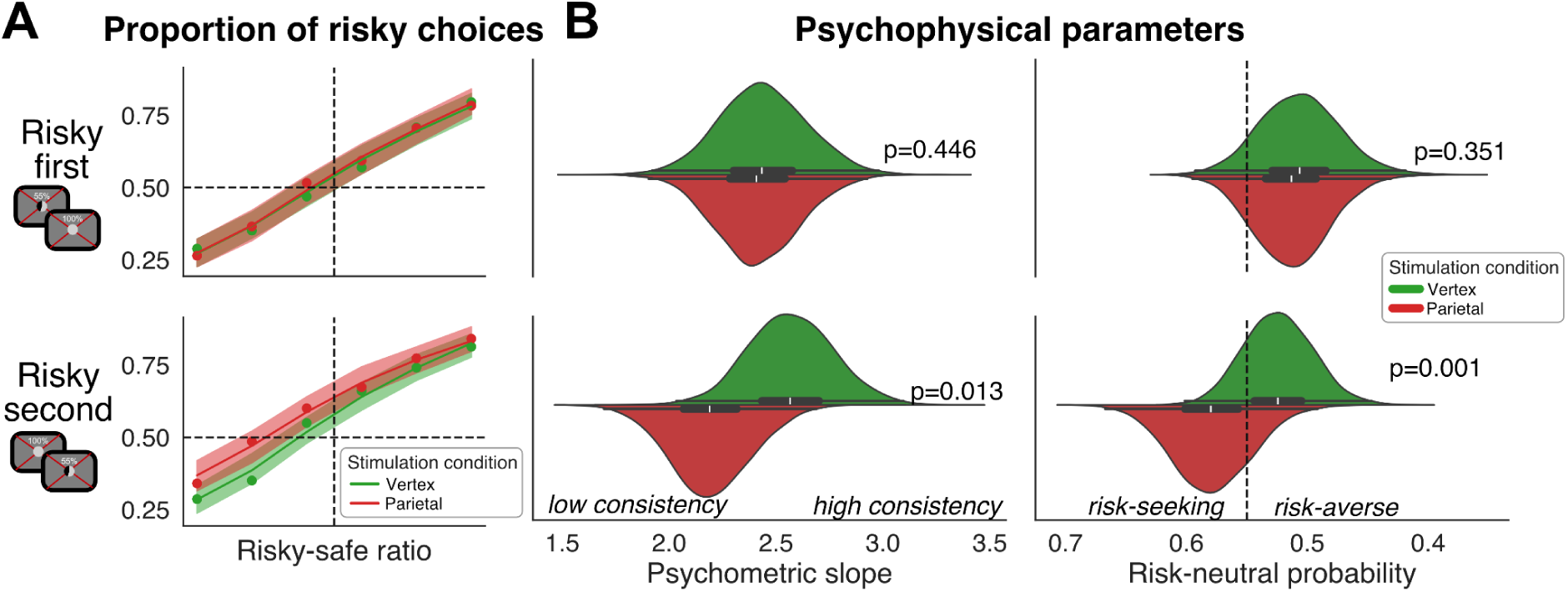
A) Psychometric Curves. Earlier work (de Hollander, Grueschow, et al., 2024) has shown a strong effect of presentation order on risk preference, in line with the assumption that first-presented options are represented more noisily in working memory and therefore more biased. Here, we replicate this effect, as participants were more likely to choose the risky option when it was presented second (compare bottom with top panel). Moreover, when risky options are presented second, subjects show a markedly more shallow response curve with a more risk-seeking indifference point (leftwards). Circles represent raw behavioral data, while the shaded areas depict the 95% credible intervals of the posterior predictions of the psychophysical choice model. The x-axis corresponds to six individually determined risky-safe ratios based on calibration data. **B) Model Parameters of psychophysical model**: In line with the qualitative effects on the psychometric slope, the estimated parameters of the psychophysical model indicate shifted risk preference towards risk-seeking and a significant decrease in response consistency after parietal TMS, but only when the risky option was presented second.

As for average risk preferences, we had no direct hypothesis about the effect of cTBS. In our Bayesian perception framework, when noise increases, subjects should rely more on their priors and become more biased in their perception of the payoff magnitudes. However, whether this leads to more or less risk averse behavior depends on multiple factors, such as the prior beliefs of the participants about the plausible ranges of payoffs, which option was presented first, and which of the two options is affected more by the increased noisiness (considering factors like heteroskedastic noise and working memory effects; de Hollander, Grueschow, et al., 2024).

Empirically, we found a very robust decrease in risk aversion after cTBS, again only when safe options were presented first, mirroring the order-specific effect of noise (risk-neutral probability increased from 52.4%, [95% CI: 47.2%, 58.3%] to 57.9% [51.8%, 64.3%], p_Bayesian_=0.001; Fig. 3B). Conversely, cTBS did not affect risk preference during trials where the risky option was presented first (risk-neutral probability increased only slightly from 50.6% [44.3%, 57.45] to 51.3% [45.3%, 57.1%], p_Bayesian_=0.351). The interaction effect between presentation order and stimulation was significant, both for the indifference point (p_Bayesian_=0.002) and choice consistency (p_Bayesian_=0.0338).

As discussed, perceptual theories of decision-making under uncertainty (Frydman & Jin, 2021; Hollander et al., 2024; Khaw et al., 2020) suggest that noisier representations of the payoff magnitudes should go hand in hand with more biased payoff percepts: The noisier the perception of a payoff magnitude, the more decision-makers should rely on their prior beliefs, thereby minimizing the average (squared) error over all possible payoffs but also increasing biases (Petzschner et al., 2015; Pouget et al., 2013). This means that decision-makers with lower choice consistency should also be more biased in their decisions (i.e., less risk-neutral). We indeed find this relationship in our experimental data, replicating earlier work (Barretto-García et al., 2023; de Hollander, Grueschow, et al., 2024; Khaw et al., 2020). Across participants, the risk preference (i.e., the indifference point of the psychophysical model) correlates highly with choice consistency (r(34)=0.76, p=0.004).

Critically, if cTBS on the right parietal cortex modulates risk preferences *via* the acuity of neurocognitive magnitude representations, one would expect that the effects of cTBS on choice consistency should also correlate with its effect on choice bias. Indeed, in our data the average decrease in choice consistency was correlated with the shift in average risk preference away from risk neutrality (r(34)=-0.51, p<0.001, one-sided). Furthermore, this correlation was slightly stronger when only considering trials where the risky option was presented second (r(34) = −0.59, p<0.001, one-sided) as compared to trials where the risky option was presented first (r(34)=-0.32, p=0.027, one-sided).

In summary, the analysis of choice behavior using psychophysical modeling shows a robust effect of parietal cTBS on both choice consistency and average risk preference, in line with the decreased fidelity of the numerically tuned signals measured using fMRI. However, the effect only occurs when safe options are presented first. We now turn to more mechanistic computational models of choice to better understand this empirical pattern.

### Shifts in risk preference are driven by increased noise in small-but not large-magnitude representations

After examining the effects of cTBS on magnitude representations in the right parietal ANS, as well as on choice consistency and average risk preferences, we applied our previously published Perceptual and Memory-based Choice (PMC) model (de Hollander, Grueschow, et al., 2024) to participants’ behavior to gain deeper mechanistic insights into the impact of cTBS on the parietal ANS. The PMC model posits that choices are guided by the maximization of perceived expected value, in contrast to traditional frameworks that emphasize objective expected value or subjective utility (Woodford, 2020). The model proposes that decision-makers perform Bayesian inference on noisy neural representations (anatomically situated in the parietal cortex), which leads to systematic ‘regressive’ biases towards the perceived mean payoff. Thus, critically, the PMC model suggests that noise in these representations can distort the perceived value of options – in particular for those options that are presented earlier, as the effects of working memory deterioration make them more susceptible to biases compared to later-presented options.

We estimated model parameters across two TMS conditions, allowing noise levels to vary for either the first option only (working memory noise) or both options (perceptual noise). While the PMC model qualitatively accounted for reduced choice consistency following parietal cTBS, it failed to capture the specific increase in risky choices when risky options were presented second (as shown in the “Weber model” in Fig. 4A). At the same time, model comparisons using leave-one-out expected log predictive density (ELPD) consistently ranked null models (no TMS effect) lowest among the six models tested (see Table 1), suggesting that cTBS has some effect that is not clearly captured by the tested version of the PMC model.

**Figure 4.**
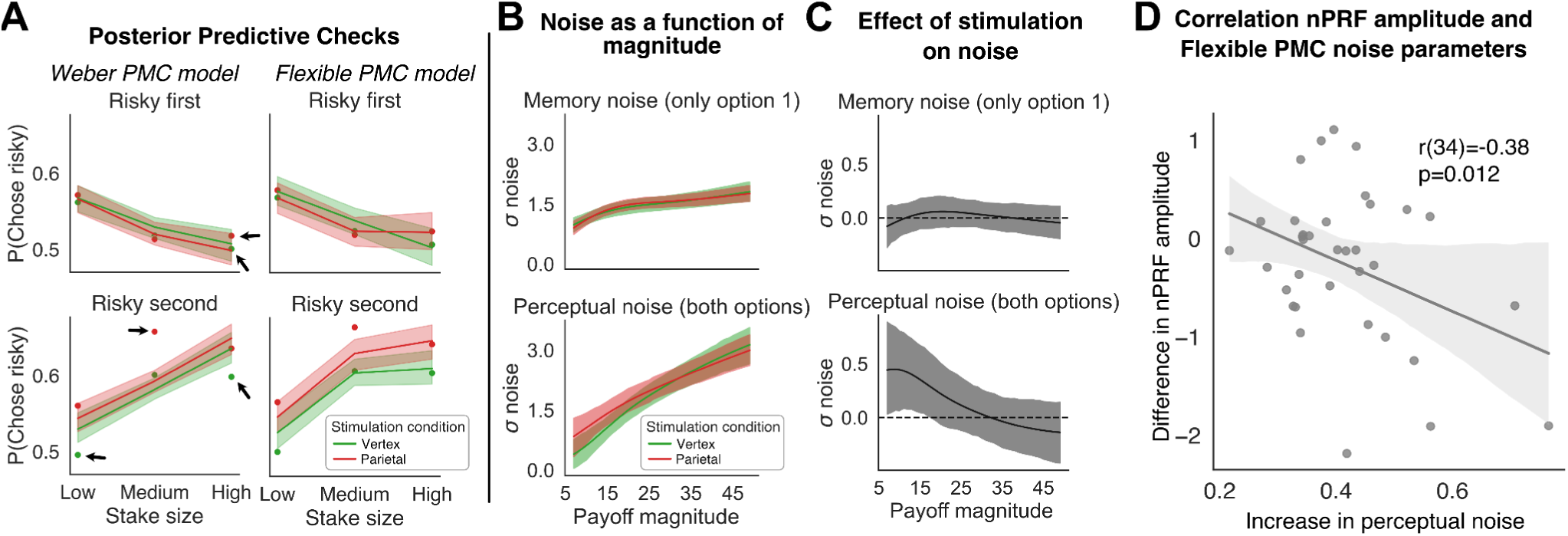
A) Posterior predictive checks: The Flexible PMC model captures the interaction between presentation order, stake, and stimulation condition on risky choices better than the Weber PMC model. In particular, the flexible PMC can capture the interaction between order and stimulation on choice proportions, unlike the simpler Weber model. The shaded areas correspond to the 95% credible interval of the group-level predictions. **B) Noise as a function of magnitude**: The model-fitted noise with which magnitudes are perceived increases markedly with the objective magnitude, in an affine trend (i.e., the noise never reaches 0, not even for small magnitudes). Conversely, working memory noise is estimated to be relatively stable across different magnitudes, although it slightly increases with magnitude as well. The shaded areas correspond to the 95% credible interval of the group-level estimates. **C) Effect of TMS on noise:** When the model-fitted noise curve of the vertex condition is subtracted from that of the cTBS condition, it becomes clear that, after parietal perturbation, noise increases for both presented options but predominantly for smaller magnitudes. The shaded areas correspond to the 95% credible interval of the group-level estimates. **D) Brain-behavior correlation:** The cTBS-induced decrease in nPRF amplitudes in the parietal cortex correlates with the cTBS-related increase in perceptual noise for magnitudes between 7 and 14.

**Table 1.**
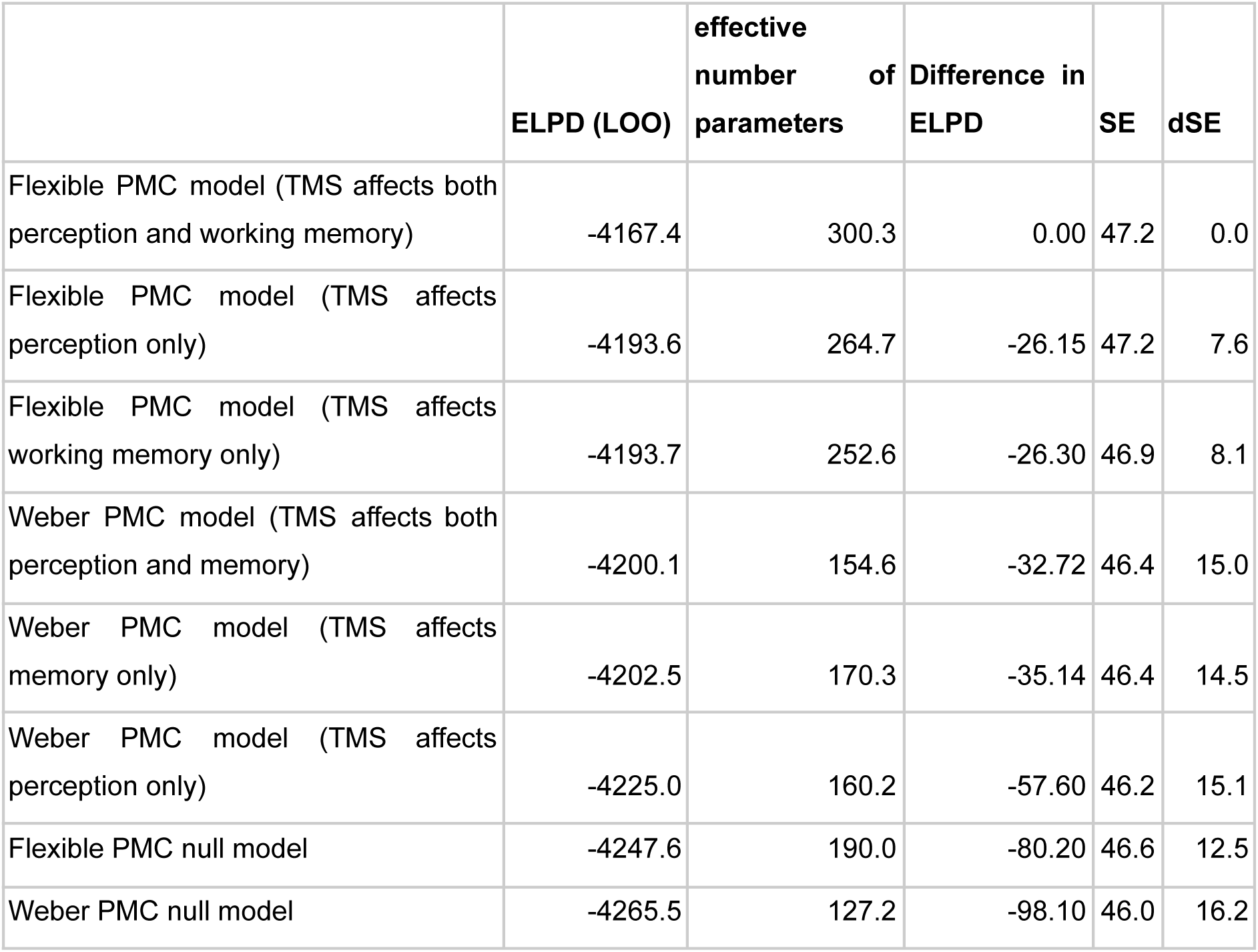
Formal model comparison using the expected leave-one-out log posterior density (ELPD; Vehtari et al., 2017) shows that the flexible noise models outperform models that assume Weber’s law. Critically, all models that assume an effect of TMS on noise outperform corresponding null models that assume no effect of TMS. Abbreviations: *ELPD (LOO)*: The expected log predictive density estimated using leave-one-out cross-validation. It measures the model’s predictive accuracy by evaluating how well the model predicts unseen data, with higher values indicating better performance. *Effective Number of Parameters*: A measure of model complexity, reflecting the number of parameters effectively used in explaining the data. Higher values suggest a more complex model. *Difference in ELPD*: The difference in expected log predictive density between a given model and the reference model. Positive values favor the given model, while negative values favor the reference model. *SE (Standard Error)*: The standard error associated with the ELPD estimate, reflecting the variability in the ELPD estimate. *dSE (Difference in Standard Error)*: The standard error of the difference in ELPD between models, representing the uncertainty in the comparison of their predictive performance (note that dSE takes into account correlations between samples across models and is therefore larger than the difference divided by mean SEs).

Since the PMC model did not fully capture the observed behavioral patterns, particularly the specific increase in risk-seeking when risky options were presented second (see the posterior predictive plots in the left panel of Fig. 4A), we followed the proposed procedure of further model refinement to better account for the observed effects (Gelman et al., 2013; Lee & Wagenmakers, 2014).

A key insight from the neural numerical PRF modeling relevant to our cognitive model development is that most numerical receptive fields identified in the fMRI data prefered smaller numerosities relative to the stimulus distribution. While the interquartile range of presented numerosities in our experimental paradigm was [13, 30], the interquartile range of preferred numerosities across all subjects was [6, 10] (see Fig. 3B). This suggests that the regions in the parietal ANS targeted by TMS primarily represent small numerosities, which may explain the misfit of our original model.

To investigate this further, we fitted a psychophysical model with a median split on average stake size (the sum of the risky and safe options). We found a significant reduction in choice consistency for low-stake trials when safe payoffs were presented first (slope from 3.02 [CI 95%: 2.73, 3.30] to 2.32 [2.05, 2.57], p_Bayesian_<0.001; see Supplementary Fig. 1), but no such reduction when risky options were presented first (from 1.73 [1.50, 2.01] to 1.59 [1.33, 1.85], p_Bayesian_=0.204). This interaction effect between stake size and stimulation condition on choice consistency was significant (p_Bayesian_=0.0153).

Thus, parietal cTBS primarily affected choices involving smaller numerosities, particularly when the safe option was presented first. This indicates that the parietal ANS region targeted in our study plays a critical role in representing and processing smaller magnitudes. Increased noise in these representations under cTBS reduces choice consistency and biases participants toward riskier options in these specific scenarios. These findings underscore the importance of considering the numerical tuning properties of the parietal ANS in understanding its role in economic decision-making.

To now capture these insights in a mechanistic computational model, we extended our PMC model by allowing noise levels to vary with magnitude, relaxing the assumption of constant noise in logarithmic space (i.e., scalar invariance; Khaw et al., 2020) and, critically, allowing us to estimate the effect of cTBS on the representational noise of different magnitude sizes. Inspired by work on magnitude averaging by Prat-Carrabin & Woodford (2022), we introduced a 5-parameter B-spline function that maps magnitudes to noisiness in natural space, on magnitude averaging, we introduced a 5-parameter B-spline function to map magnitudes to noisiness in natural space, ensuring a non-linear yet smooth relationship (see methods).

To make sure that this new *Flexible PMC model* was not overly complex (’overfitting’ the data), we pitted it against the original *Weber PMC model* using both qualitative and quantitative model comparison. Qualitatively, the Flexible PMC model explained the order-specific increase in risky choices far better than the original Weber PMC model (Fig. 4A). Furthermore, formal model comparison revealed that the best-fitting model included both memory-specific noise (for the first option) and shared perceptual noise, both modulated by TMS. This model consistently outperformed all other variants, including a null model (Table 1), confirming that the effect of TMS is robust and restricted to a specific range of payoff magnitudes. Parameter estimates further highlight that it was the shared perceptual noise for smaller payoff magnitudes (approximately 7–14) that differed most significantly between TMS conditions, with credible intervals excluding 0.0 (Fig. 4B and C). These findings provide a refined mechanistic account of how parietal ANS disruptions influence decision-making under uncertainty.

### Linking Neural and Behavioral TMS Effects

Our findings reveal that cTBS of the parietal ANS disrupts neurocognitive magnitude representations, leading to reduced choice consistency and increased risk-seeking behavior. These effects were particularly pronounced for smaller numerosities and when safe options were presented first. A refined “Flexible PMC model” that accounts for magnitude-dependent noise captures this pattern, by showing that cTBS specifically modulated perceptual noise for smaller magnitudes. This aligned with evidence from the fMRI analyses that numerical tuning in the parietal cortex is primarily focussed on smaller numbers (Barretto-García et al., 2023; Cai et al., 2021; Harvey et al., 2013).

A key prediction of our theoretical framework, which posits the parietal ANS as the neural substrate for the neurocognitive representation of numbers, is that disruptions in parietal numerical representations should lead to increased neurocognitive noise, resulting in corresponding behavioral changes. To test this comprehensively, we examined whether the decrease in numerical receptive field (nPRF) amplitude in the parietal cortex predicted the increase in noise for small numerosities. Consistent with this hypothesis, we found a significant correlation (r(34) = 0.38, p = 0.012, one-sided; Fig. 4D) between the relative decrease in nPRF amplitude in numerically tuned voxels and the average increase in neurocognitive noise across the lowest three safe options (7/10/14) according to the flexible PMC model.

Taken toegether, our results provide strong evidence that disruption of parietal ANS representations of smaller magnitudes increase perceptual noise for smaller payoffs, reducing choice consistency and altering risk preferences. The alignment of behavioral and neural effects under TMS supports our causal framework, highlighting the critical role of parietal magnitude representations in economic decision-making. This integrated neural-cognitive approach explains how disruptions in numerical representations lead to systematic behavioral biases, offering a mechanistic account of individual differences in decision-making under uncertainty.

## Discussion

Everyday decisions often involve choices between more and less uncertain outcomes, and individuals differ in their preferences for how to handle such choices. Recent models and behavioral findings suggest that such preferences may not only reflect subjective valuation processes, but perhaps also be the precision and distortions with which individuals perceive and represent the relevant numerical magnitudes in such tasks, such as the potential payoffs of different options, before any decision is made (Barretto-García et al., 2023; de Hollander, Grueschow, et al., 2024; Frydman & Jin, 2021; Khaw et al., 2020). However, from a neural point of view, even though some neuroimaging studies have revealed correlational links between risk preferences and the fidelity of neural signals in the ‘approximate number system’ (ANS) in the parietal cortex (Bueti & Walsh, 2009; Dehaene, 2011; Walsh, 2003)(Barretto-García et al., 2023; de Hollander, Grueschow, et al., 2024) a direct causal link between the precision of magnitude representations in the parietal ANS and economic decision-making yet has to be established. In this study, we establish such a link, by perturbing parietal ANS function using continuous theta-burst rTMS (cTBS). Importantly, we employed a novel approach using numerical receptive field modeling (nPRF) on individual fMRI data to precisely target regions of the dorsal parietal cortex with robust tuning for payoff magnitudes in each participant. This innovative use of nPRF modeling allowed us to probe the mechanistic effects of cTBS on parietal magnitude representations with unprecedented precision. We found that the amplitude of numerically tuned responses decreased after cTBS, while their tuning profile stayed constant. As a result, the amount of information about presented numerosities that could be decoded using an inverted nPRF model was reduced. Notably, the targeted area predominantly contained preferred responses to smaller numerical magnitudes. This neural preference for smaller magnitudes was reflected in the behavioral effects of parietal cTBS, as choice behavior was modulated only during decisions involving relatively small stake sizes.

To achieve a more detailed mechanistic understanding of these effects, we applied computational cognitive modeling. Specifically, we extended our existing Perceptual and Memory-based model of risky Choice (PMC; de Hollander, Grueschow, et al., 2024), which frames simple choices between prospects as a problem of Bayesian inference on noisy neurocognitive representations. While the original model assumed scalar invariance—a strictly linear increase in noise with increasing magnitudes—we developed a novel version that relaxes this assumption. Building on earlier work (Prat-Carrabin & Woodford, 2022), the updated model employs flexible spline functions to estimate the relationship between magnitude and cognitive noise based on observed data. Although this added flexibility increased the model’s complexity, qualitative and formal comparisons confirmed that it was necessary to capture the intricate empirical patterns. More importantly, parameter estimates revealed that cTBS specifically increased the noisiness of representations for smaller payoffs, leading to perceptual distortions that accounted for the observed rise in risk-seeking behavior in a specific subset of choices.

An earlier, related study that applied brain stimulation in the context of decision-making by Coutlee et al. (2016) showed that 1-Hz offline rTMS applied to the left parietal ANS (led to decreased risk-taking. In contrast, our study observed increased risk-taking following offline theta burst stimulation of the right IPS. An important difference between the two sets of results is that Coutlee et al. did not separately examine the effects of TMS on choice consistency or average preferences. However, the authors do report that after IPS stimulation, participants were not only less risk-seeking but also less likely to choose options with a higher expected value—a hallmark of both decreased choice consistency and increased bias. This additional finding can reconcile the diverging results of raw choice proportions. Such a reinterpretation of earlier results in the literature highlights the importance of jointly considering the effect of experimental manipulations on both average risk preference (i.e., an inferred indifference point or utility function), as well as choice consistency (Olschewski et al., 2022; Olschewski & Rieskamp, 2021).

Besides these conceptual considerations, four potentially relevant methodological differences with the Coutlee et al. study should also be briefly mentioned. First, Coutlee et al. stimulated left IPS rather than right IPS (and there is some evidence for lateralizing risk-taking behavior; Dantas et al., 2023; Sacré et al., 2019). Note that we chose to stimulate right IPS based on literature in numerical cognition that suggest right-lateralization of the ANS (Barretto-García et al., 2023; Dehaene, 2011; Dehaene et al., 2003; Eger et al., 2003; Lasne et al., 2019), as well as our work that showed the most robust links between neural activity and choices in right parietal cortex (Barretto-García et al., 2023; de Hollander, Grueschow, et al., 2024). Second, Coutlee et al. used a different stimulation protocol: 1-Hz offline repetitive TMS rather than theta-burst stimulation. Third, they had a decision-making task that involved symbolic rather than non-symbolic magnitudes. Fourth, their task had no explicit working memory component.

A similar study by Panidi et al. (2023) also showed that stimulation of the left and right “posterior parietal cortex” using cTBS led to increased risk aversion in a multiple price lists task, where a set of multiple gambles with increasing payoff are presented together versus a baseline option. The increase in risk aversion is consistent with an increase in neurocognitive noise (and increased bias), just as observed in our study. However, the authors also report increased choice consistency as estimated by a prospect theory model. Unfortunately, the choice consistency derived from prospect theory is hard to compare to our results because it is defined based on the subjective scale of the estimated utility function, rather than the objective scale of actual payoffs we use here. Moreover, the multiple price lists-paradigm comes with severe methodological problems (Drichoutis & Lusk, 2016) and might elicit additional cognitive processes over and above numerical processing, such as flexible spatial attention, which could have been influenced by cTBS as well.

Other relevant work applying brain stimulation techniques (Dantas et al., 2021, 2023) revealed that risk preference can also be modified by stimulation of the frontal cortex. Specifically, risk appetite has been manipulated by increasing or decreasing theta band power in frontal areas using transcranial alternating current stimulation (tACS), with opposite effects for the two hemispheres. The authors interpret this finding as an increase/decrease of cognitive control, implemented by frontal oscillations and helping a participant “inhibit” default responses (see also Gianotti et al., 2008). A potential alternative explanation, based on our perceptual modeling framework, could be that frontal theta oscillations may increase selective attention toward relevant stimuli (Landau et al., 2015; Spyropoulos et al., 2018). Upregulated selective attention might improve magnitude perception and thereby shift risk appetite. Future work should explore this hypothesis by explicitly measuring noise in both choice and parietal ANS after frontal brain stimulation. Interestingly, the processing of symbolic versus non-symbolic numerosities is somewhat lateralized as well (left- versus right-hemisphere respectively; Dehaene & Cohen, 1991; Warrington & James, 1967), which might explain why theta band oscillations in left hemisphere are related to risk appetite symbolic risk tasks (Dantas et al., 2021, 2023) and right-hemispheric theta oscillations are related to non-symbolic risk tasks (Gianotti et al., 2008).

Although neuroimaging studies on the parietal ANS and risk-taking have shown a link between numerical magnitude processing and the parietal cortex (Barretto-García et al., 2023; de Hollander, Grueschow, et al., 2024) and brain stimulation studies also hinted to a causal role of the parietal cortex in risky choice (Coutlee et al., 2016; Panidi et al., 2023), a direct causal link between the precision of parietal magnitude representations and behavior was still lacking. By combining brain stimulation (cTBS) and neuroimaging (fMRI) techniques, our study provides compelling evidence for a critical causal role of the perceptual processing of magnitude information in economic choice problems and the right parietal ANS in risky decision-making.

Traditionally, studies on decision-making involving risks have mainly focused on the classical ‘valuation network’ including the ventromedial prefrontal cortex, as well as the insula and the dopaminergic system, including the dopaminergic midbrain and the ventral striatum (Bartra et al., 2013; Mohr et al., 2010). This focus on the valuation network has also driven the search for clinical markers of psychiatric disorders like gambling addiction (Miedl et al., 2012; Sharp et al., 2012). However, our findings suggest that neural systems involved in perceiving decision-making problems might play a similarly significant role in decision-making under uncertainty. That is, some behavioral research already suggests that the expectations about potential payoffs are structurally different in gamblers (Griffiths, 1990; Rogers, 1998; Spurrier & Blaszczynski, 2014). Moreover, structural differences in perceptual brain regions regions, particularly the right parietal ANS, could contribute to psychiatric disorders and warrant further investigation. Notably, previous studies have demonstrated that structural properties of the right posterior parietal cortex predict individual differences in risk preferences among healthy adults (Gilaie-Dotan et al., 2014). Future work might investigate false beliefs in pathological gamblers using explicit Bayesian models of risk perception (Paliwal et al., 2014, 2019) and link those to parietal regions involved in numerical cognition.

Our findings reveal that brain state can have a profound influence on economic choices (see also Chew et al., 2019; de Hollander, Grueschow, et al., 2024). This has important implications for risk elicitation techniques used in real-world applications, such as in the finance industry (Alserda et al., 2019), and might also further elucidate the relationship between arousal/attentional states and gambling behavior (Brown, 1986; FeldmanHall et al., 2016). Understanding how arousal and attentional states modulate magnitude representations could also shed light on the mechanisms underlying gambling behavior and other risky activities, potentially offering novel therapeutic targets for interventions. Future research should explore how these dynamic neural and cognitive factors interact to shape economic behavior, paving the way for more personalized approaches to managing risk and decision-making in both clinical and everyday contexts.

Both our neuroimaging and behavioral results following cTBS suggest that relatively smaller—rather than larger—numerical magnitudes are specifically encoded in the parietal cortex. It remains an open question whether parietal neurons genuinely specialize in encoding smaller magnitudes, or if this finding reflects a potential property of our experimental design in which smaller magnitudes were more frequent. This design choice could have facilitated responses to smaller numbers, or could have introduced range adaptation effects. Future studies that systematically vary the range of potential payoffs will be crucial for disentangling these hypotheses (Cai et al., 2021). Regardless, our findings underscore the utility of focal brain stimulation techniques like TMS, particularly when guided by precise fMRI-based functional mapping, in decoding neural representations of abstract stimulus features such as numerosity or magnitude. By directly manipulating these neural coding patterns, TMS provides a powerful tool to establish causal links between neural activity and cognitive functions.

Advances in ultra-high field neuroimaging (Dumoulin et al., 2018), with its considerably improved resolution and signal-to-noise-ratio, combined with dense individual sampling across multiple sessions (Poldrack, 2017), should enable even more precise targeting of numerosity-tuned regions in the parietal cortex in future work. Such an approach could go beyond general targeting of the ANS and towards selectively stimulating cortical patches corresponding to specific segments of the number line within the topographically organized parietal numerosity map. Such an approach could serve as a numerical analog to phosphene mapping in the primary visual cortex using TMS (Murphey, 2009). This could potentially tighten the gap between observed neural coding patterns and behavior in healthy human participants even further.

In conclusion, our brain stimulation study provides compelling evidence for a critical causal role of the right parietal approximate number system in risky decision-making. Moreover, it offers a detailed, mechanistic neural account of the parietal ANS’s involvement in magnitude representation through nPRF models and how perturbations to this region affect magnitude perception and, consequently, risk preferences. These findings highlight the integral role of perceptual processes in tasks traditionally considered purely economic. Building on this perspective, our findings underscore the need for further development of economic decision-making models that integrate both perceptual and preference-based components (Woodford, 2020) and demonstrate how advanced neuroimaging and brain stimulation approaches can push theoretical frameworks beyond what is achievable through behavioral experiments alone.

## Methods

### Participants

Seventy-eight right-handed participants (31 females, ages 18 to 35; mean age 23.9) volunteered to participate in this study. We informed them about the study’s objectives, the equipment used in the experiment, the data recorded and obtained from them, the tasks involved, and their expected payoffs. We also screened participants for MRI and TMS compatibility before they participated in the study. No participant had indications of psychiatric or neurological disorders or needed visual correction. Our experiments conformed to the Declaration of Helsinki, and our protocol was approved by the Canton of Zurich’s Ethics Committee. After the first sessions, thirty-five participants (12 females; age 18 to 30; mean age 23.1) were selected for two more follow-up experimental sessions (see below for more details about the procedure).

### General procedure

All 78 participants attended an initial experimental session, during which they received an informed consent form and detailed task instructions. After providing consent, participants were familiarized with the TMS system, and their active motor threshold was determined in the left primary motor cortex (M1; see below). Following this small TMS session, participants proceeded to the scanner room, where structural MRI data were collected. During the acquisition of the structural MRI, participants performed a calibration task to estimate their average risk preference and choice consistency.

This calibration task involved making choices between 96 risky gambles (55% chance of payout and 45% chance of winning nothing) and 96 corresponding ‘safe options’ (sure payout). Each choice pertained to one of 5 unique safe payoffs (7, 10, 14, 20, or 28 CHF; 1 CHF is currently 1.12 USD or 1.07 EUR, Big Mac index ∼= 0.154) and a risky option that was 2^ℎ/4^ times the safe option, with ℎ all integers from 1 to 8. Each pair was presented twice, once with the safe option shown first and once with the risky option shown first. Calibration data were fitted using a psychometric probit model, with ‘chose risky option’ as the dependent variable and an intercept and the log-ratio of the risky and safe options as independent variables (de Hollander, Grueschow, et al., 2024). The fitted model predicted, for any given risky/safe payoff ratio, a proportion of risky choices.

Based on these predictions, a participant-specific task design was developed to maximize the precision of psychophysical model estimation (de Hollander, Grueschow, et al., 2024; Heerema et al., 2023). This design used six payoff fractions spaced equally in log-space, corresponding to predicted risky choice proportions between 20% and 80%. These were combined with five safe payoffs (7, 10, 14, 20, 28), with each combination presented twice in both safe-first and risky-first orders, resulting in 120 trials per session. After calibration, this task design was presented during fMRI scanning over six runs of 20 trials each (see Experimental Paradigm).

Following the first session, participants were monetarily compensated, and their data were analyzed to assess suitability for TMS sessions. Suitability was determined based on four criteria: (1) a robust cluster of at least five voxels in the dorsal parietal cortex showing numerosity tuning with at least 10% explained variance (R2R^2R2) across all trials; (2) anatomical accessibility of the numerosity-tuned region for TMS (e.g., on the dorsal bank of the IPS); and (3) no adverse effects of TMS pulses on M1 (4) no extreme choice behavior in the task (less than 10% or more than 90% risky choices across trials).

Thiry-two (32) subjects were excluded because they did not show a robust numerically-tuned cluster in IPS, 5 subjects were excluded because the found cluster lay too deep int he sulcus to be reached by TMS, 3 subjects showed adverse effects of TMS, one subject excluded herself and two subjects showed too extreme choice behavior.

Thus, out of 78 participants, 35 met the inclusion criteria and were invited for two additional sessions. The relatively low number of eligible participants compared to earlier studies (Barretto-García et al., 2023; de Hollander, Grueschow, et al., 2024) was largely due to the reduced number of trials in this study (120 compared to 192 and 216 in prior studies). This choice was based on evidence that cTBS influences behavior for approximately 30 minutes (Huang et al., 2005), and we aimed to select participants whose numerosity-tuned responses were sufficiently reliable to observe within this time window.

It is important to note that our study employed a within-subject design, ensuring that baseline differences between participants do not confound TMS effects. While participants selected for TMS exhibited slightly higher choice consistency (2.25 [CI: 1.92–2.56] vs. 1.94 [CI: 1.65–2.24]) and were slightly more risk-seeking (neutral risk probability: 48.3% [42.9–54.3] vs. 44.7% [38.5–51.1]), these differences were not statistically significant (p_Bayesian_=0.082) and do not reflect a selection based on risk preferences. Instead, the selection was strictly guided to ensure accurate anatomical targeting of numerosity-tuned regions. The use of individualized targeting was critical due to the high interindividual variability in the location of numerosity-tuned populations (Harvey et al., 2013, 2020). This approach is significantly more effective than targeting standardized coordinates (e.g., MNI152 or fsaverage) and cannot explain away the observed within-subject effects of TMS (Sack et al., 2009).

In the two experimental sessions, participants got to read the instructions for the task again and gave written informed consent. Then, they immediately went to the scanner room, and the active motor threshold was determined using the TMS device in the scanner room (see TMS stimulation for details). Then, the participant laid down on the scanner bed and was stimulated using the cTBS protocol, targeting either the vertex or the predetermined parietal cluster. After the stimulation protocol finished, the participant was immediately put in the scanner, and the experimental paradigm was started as soon as possible. The average time that elapsed between the end of the cTBS protocol and the onset of the task was 3:12 (std. 0:19). Out of 35 participants, 18 participants were stimulated over the parietal cortex for the 2nd session and stimulated over the vertex for the 3rd session. For the remaining 17 participants, this order was reversed.

### Experimental paradigm

The task of the participants was, for every trial, to choose between a certain amount of money (7/10/14/20 or 28 CHF) and a gamble with a 55% probability of winning a larger amount of money and a 45% probability of winning nothing at all. The choices were represented by a sequence of tailored stimuli. The general sequence is illustrated in Fig. 1A. The screen always contained a red cross with two diagonal lines to keep fixation near the center of the screen and to not confound the numerosity of stimuli like a standard fixation cross or point might (Harvey et al., 2013). The start of a new trial was indicated by the fixation cross turning green for 250 ms. Then, after a pause of 300 ms with just a red fixation cross, a pile chart with a diameter of 1 degree-of-visual-angle (dova) was presented for 300 ms to indicate the probability-of-payout for the coming stimulus. This was always either 55% or 100%. Then, after another 500 ms of just fixation cross, a stimulus array of 1-CHF coins appeared that represented the potential payoff of the first choice option. The coins had a radius of 0.3 dova and were all randomly positioned with their centre within a circular aperture with a diameter of 5.25 dova. The coin stimulus array was presented for 600 ms, after which only the fixation cross was presented for a jittered duration of either 5, 6, 7, or 8 seconds. Then, a pie chart indicating the probability of the second payoff was presented for 300 ms, followed by a 300ms fixation screen, and another coin stimulus array, representing the potential payoff of the second option, again for 600 ms. As soon as the second stimulus array was presented, participants could indicate their response with their index (first-presented option) or middle finger (second-presented option). As soon as they responded, they saw a 1 or 2 indicating which option they had chosen for 500 ms. After the coin stimulus pile was presented, the remaining duration of the trial was either 4, 4.5, 5, or 5.5 seconds.

Stimuli were presented using a projector on a screen at the back of the bore and a hot mirror system on top of the coil system (1920 x 1080). The projector screen was 125cm from the participant’s eyes and was 42 cm wide, corresponding to a field-of-view of approximately 19 dova.

### TMS stimulation

At the start of session 2 and 3, the motor hotspot of the left M1 was identified by locating the point eliciting the strongest movement-evoked potentials (MEPs) in the first dorsal interosseous (FDI) muscle using a figure-of-eight TMS coil. A circular grid was mapped onto each participant’s anatomical MRI using a neuronavigation system (BrainSight, Rogue Research Inc., Canada), focusing on the hand motor region in the anterior central sulcus. The motor hotspot was determined as the grid point producing the strongest FDI MEPs. Participants then performed an active motor task (pressing thumb and index finger with ∼20% maximum force) to determine the active motor threshold (AMT), defined as the lowest TMS intensity eliciting MEPs ≥ 200 μV in 5 out of 10 consecutive pulses. The AMT was retested visually in the scanner room by observing FDI twitches. The mean AMT was 50.8% (SD = 7.9%) outside the scanner and 54.5% (SD = 6.8%) inside the scanner room.

During combined TMS-fMRI sessions, participants received cTBS by means of a Magenture MagPro X100 stimulator (Magventure A\S, Farum, Denmark) targeting either the individual numerosity population receptive field (nPRF) cluster in the intraparietal sulcus (IPS) or the vertex, determined using neuronavigation. The standard cTBS protocol (Huang et al., 2005) involved bursts of three 50 Hz stimuli repeated at 5 Hz for 40 seconds, delivering 600 pulses at 80% of the participant’s AMT. The coil (MRi-B91, Magventure) was positioned tangentially to the cortical surface, with the handle pointing posteriorly, and stimulation coordinates were marked on a latex cap affixed to the participant’s head. The vertex control site was defined as the intersection of the left- and right-central sulci in the interhemispheric fissure.

### Cognitive computational modeling

#### Psychophysical (probit) model

In line with earlier work (Barretto-García et al., 2023; de Hollander, Grueschow, et al., 2024; Khaw et al., 2020; Olschewski et al., 2022; Olschewski & Rieskamp, 2021), we modeled the risky choice data using a standard psychophysical function, which models choice consistency and average preference as two distinct latent cognitive variables. Note that the psychophysical model we use here is a *measurement model* and thus remains agnostic to the precise underlying mechanisms. For example, the noise in choices could be due to noise in valuation or perception (Barretto-García et al., 2023).

Specifically, the psychophysical can be parameterized as a generalized linear model:

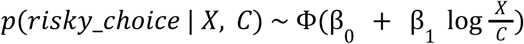

Here, 𝑋 is the potential monetary payoff for the risky option, 𝐶 is the potential payoff for the safe option, and Φ is the standard cumulative probability function of the standard normal distribution (i.e., probit function). β_0_ and β_1_ are free parameters that are estimated. Together, they quantify (a) the choice consistency (slope β_1_) and (b) the indifference point of the decision-maker (here we use the the risk-neutral probability 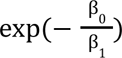. The psychophysical function was estimated using Bayesian estimation and as a hierarchical generalized linear model as implemented in the *bambi* library (v 0.13; Capretto et al., 2022). See de Hollander et al. (2024) for more details on the links between psychophysical models and perceptual theories of risky choice (e.g., Khaw et al., 2020).

#### The Perceptual and Memory-based model of risky Choice (PMC model)

We will now briefly discuss our Perceptual and Memory-based model (PMC) of risky choice. For a more complete introduction, see de Hollander et al. (de Hollander, Grueschow, et al., 2024).

Briefly, the PMC model assumes that decision-makers try to maximize *expected payoff*, based on *noisy representations* of the choice problem (see also Khaw et al., 2020). More formally, when participants decide on a risky option with a payoff 𝑋 (and a probability of payout 𝑝) and a safe option with a payoff of 𝐶, they only have access to two corresponding random variables 𝑟_𝑥_ and 𝑟_𝑐_ (their ‘neurocognitive representations’) that are conditional on 𝑋/𝐶:

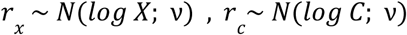

Crucially, to optimize their expected value in the long run, decision-makers take into account any prior beliefs they have about potential payoffs:

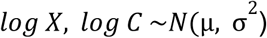

So, for a given 𝑟_𝑥_ or 𝑟_𝑐_, the participant’s estimate is a weighted sum of the center of the likelihood and the center of the prior, weighted by their relative dispersion β:

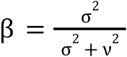

Specifically, the conditional expected values of the two options conditional on 𝑟 are:

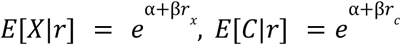

where α is a function of the mean and standard deviation of the prior:

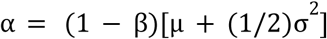

(Rational) participants will choose the risky option if and only if 𝑝 ×𝐸[𝑋|𝑟] > 𝐸[𝐶|𝑟] or, equivalently, log 𝑝 + β𝑟_𝑥_ > β𝑟_𝑐_, or log 𝑝 + β𝑟_𝑥_ − β𝑟_𝑐_ > 0 Since both 𝑟_𝑥_ and 𝑟*_c_* are (independent) Gaussian random variables, conditional on 𝑋 and 𝐶, their difference is a Gaussian variable as well:

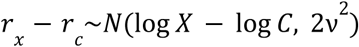

Crucially, this means that the probability that the participant chooses the risky option (log 𝑝 + β𝑟_𝑥_ − β𝑟_𝑐_ > 0) is conditional on 𝐶 and 𝑋 can be described using the cumulative normal distribution Φ(𝑥):

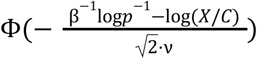

Now, the more extended PMC model adds two model mechanisms to this derivation: Working memory effects and different priors for risky versus safe options.

##### 1. Working memory effects

Options that are presented earlier in time could be more noisy. Thus, rather than a single noise parameter ν, the PM model contains two noise parameters: ν_1_ for the option that is presented first and ν_2_ for the option that is presented second. Pilot work has shown that ν_1_ and ν_2_ are correlated aross participants, which can hinder estimation. Moreover, it seems plausible that two latent traits underlie the noisiness of the two options: 1) general numerical perception noise that drives both, and 2) working memory noise. Thus, to aid parameter estimation and improve model interpretability, we reparameterize the model as follows:

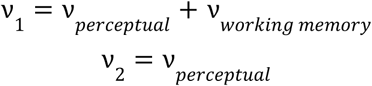

##### 2. Different priors

Risky options have, on average, considerably higher potential payoffs than safe options. For example, in the data described here, the mean safe payoff was 15.82 CHF (std. 7.50), whereas the mean risky payoff was 36.17 (std. 22.1). Potentially, decision-makers could take this into account, and this should improve their expected payoff in the long run (and earlier work has shown strong evidence for this; de Hollander, Grueschow, et al., 2024). Therefore, the PMC model includes two sets of two parameters describing the priors for risky and safe options: µ_𝑟𝑖𝑠𝑘𝑦_, σ_𝑟𝑖𝑠𝑘𝑦_ and µ_𝑠𝑎𝑓𝑒_ and µ_𝑠𝑎𝑓𝑒_.

#### PMC model summary

To summarize, the full PMC model as proposed by de Hollander et al. has the following six parameters:

1. ν_𝑝𝑒𝑟𝑐𝑒𝑝𝑡𝑢𝑎𝑙_ the amount of noise that is inherent in number perception and pertains both to numerical magnitude kept in working memory (i.e., n1) and magnitudes that are more directly accessible via a stimulus (i.e., n2).
2. ν_𝑤𝑜𝑟𝑘𝑖𝑛𝑔 𝑚𝑒𝑚𝑜𝑟𝑦_ the amount of noise that gets added to ν _𝑝𝑒𝑟𝑐𝑒𝑝𝑡𝑢𝑎𝑙_ when a magnitude has to be kept in working memory.
3. µ_𝑟𝑖𝑠𝑘𝑦_ the mean of the prior belief over risky options.
4. σ_𝑟𝑖𝑠𝑘𝑦_ the standard deviation of the prior belief over risky options.
5. µ_𝑠𝑎𝑓𝑒_ the mean of the prior belief over safe options.
6. σ_𝑠𝑎𝑓𝑒_ the standard deviation of the prior belief over safe options.

Note that although the number of parameters is relatively high, they are strictly necessary to explain the strong interaction between order and stake size on risky choice proportions that was clearly present in both datasets described in de Hollander et al. (2024), as well as in the dataset described in this paper. Furthermore, in line with these qualitative patterns, formal model comparisons on all three datasets confirm that the increased complexity of the PMC model compared to simpler model was warranted (Khaw et al., 2020).

#### The flexible PMC model

As described in the main text, the original PMC model can only model the impact of cTBS stimulation on all perceptual noise, either on both options or only on the first/second option. However, since only small numerosities can be visualized in the parietal ANS, it is plausible that only the neurocognitive representation of smaller numerical magnitudes were perturbed by parietal cTBS. Furthermore, recent evidence suggests that the assumption of Weber’s law on numerical magnitude representations might be too strict (Prat-Carrabin & Gershman, 2024; Prat-Carrabin & Woodford, 2022).

Therefore, inspired by the work of Prat-Carrabin & Woodford (Prat-Carrabin & Woodford, 2022), who used low-order polynomials to describe a non-linear mapping between numerical averages and perpetual noise, we reformulated the PMC model in such a way that the perceptual/working memory is no longer estimated as a single noise parameter, but rather as a non-linear smooth function. This allows for specific effects of cTBS on specific parts of the number line. To ease and regularize parameter estimation, we used spline regression. Concretely, the noise for a specific numerical magnitude is modeled as a linear combination of 𝑚 splines:

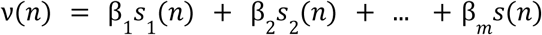

where 𝑠_1_ is the 1st 3rd degree spline, with bounds between 7 and 112 and β_1_ … β_𝑚_ are free parameters that are estimated. After some initial model comparisons, we chose to use 5 splines for our purposes. Our implementation in Braincoder uses the widely used *Patsy*-package (Gates, 2023) to define the spline functions.

More specifically, in the case of the PMC, two flexible noise functions are estimated, one for perceptual noise:

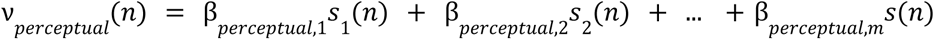

and one for working memory noise:

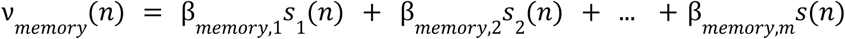

This flexible version of the PMC model has a substantially higher number of parameters than the original PMC model. Note, however, that the original PMC model can not explain the empirical behavioral patterns observed after parietal cTBS (i.e., increased risk-seeking but only for specific presentation orders). Moreover, we used a state-of-the-art formal model comparison technique, Expected Pointwise Likelihood Densities (Vehtari et al., 2017), to make sure that we did not overfit the data and the increased complexity of the model was warranted.

#### Model estimation

The PMC and Flexible PMC model are implemented in our Python toolbox *bauer* (de Hollander, Renkert, et al., 2024), which incorporates a variety of psychophysical and Bayesian decision models. They are estimated using Bayesian hierarchical estimation, built upon the computational graph framework of *pymc* (Patil et al., 2010), which uses the state-of-the art Hammiltonian MCMC sampler NUTS (Hoffman & Gelman, 2011).

A key design choice was to model individual parameters using an ‘offset’-parameterisation. The ‘offset’ parameterization guards against a funnel in parameter space by re-expressing individual-level parameters in terms of deviations from the group mean. In traditional hierarchical models, directly estimating individual parameters can result in poor identifiability when the group-level standard deviation σ is small. This creates a “funnel” shape in the posterior distribution, where individual parameters are tightly constrained near the mean, leading to high curvature and inefficient sampling. Thus, we use the following definition:

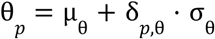

Where θ_𝑝_ is a participant-specific parameter (e.g., ν_𝑝𝑒𝑟𝑐𝑒𝑝𝑡𝑢𝑎𝑙_ or µ_𝑟𝑖𝑠𝑘𝑦_ for participant 𝑝) that is not directly estimated, but defined as a combination of (a) µ_θ_ – the group mean for that parameter, (b) σ_θ_ is the standard deviation of the group distribution, and (c) the participant-specific parameter δ_𝑝,θ_. Here, δ_𝑝,θ_ are unconstrained, and the hierarchical structure separates the scale (group-levelσ_θ_) from individual differences. This improves sampling efficiency and mitigates the funnel effect by allowing the model to explore the posterior more effectively, even when σ_θ_ is small.

To model the experimental effects of session and TMS, we use a random effects approach, where each participant has a design matrix that defines a specific parameter value on a specific trial, based on a linear combination of intercept- and condition-wise estimates:

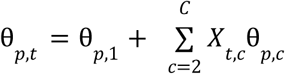

Here, θ_𝑝,𝑐_ is the parameter value for participant 𝑝 on trial 𝑡, θ_𝑝,1_ is the participant-specific intercept, 𝑋_𝑡,𝑐_ is the design matrix encoding the experimental conditions for trial 𝑡 (starting with the second column for condition effects), and θ_𝑝,𝑐_ are the participant-specificcondition-wise effects for 𝑐 = 2, … 𝐶.

The random effects approach allows both the intercept (θ_𝑝,1_) and condition-specific effects (θ_𝑝,𝑐_) to vary across participants, capturing individual differences in their responses to the experimental manipulations. However, each participant-specific effect θ_𝑝,𝑐_ is drawn from a group-level distribution:

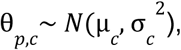

where µ_𝑐_ is the group-level mean for condition 𝑐, and σ_𝑐_^2^ is the group-level variance.

This hierarchical structure accounts for between-participant variability while enabling inference about group-level effects. By regularizing participant-specific deviations through the group-level priors, this approach avoids overfitting, ensures stable parameter estimation, and enhances the sensitivity to detect session- and TMS-related effects.

All the models and their implementation are freely accessible at https://github.com/ruffgroup/bauer/tree/main/bauer.

### MRI scanning parameters

We acquired functional MRI data using the Philips Achieva 3T whole-body MR scanner equipped with a 32-channel MR head coil, located at the Laboratory for Social and Neural Systems Research (SNS-Lab) of the UZH Zurich Center for Neuroeconomics. In all three session, we collected 6 runs of fMRI data with a T2*-weighted gradient-recalled echo-planar imaging (GR-EPI) sequence (130 volumes + 5 dummies; flip angle 90 degrees; TR = 2286 ms, TE = 30ms; matrix size 96 × 70, FOV 240 × 175mm; in-plane resolution of 2.5 mm; 39 slices with thickness of 2.5 mm and a slice gap of 0.5mm; SENSE acceleration in phase-encoding direction (left-right) with factor 1.5; time-of-acquisition 5:05 minutes). During the first scanning session, we acquired high-resolution T1-weighted 3D MPRAGE image (FOV: 256 × 256 × 170 mm; resolution 1 mm isotropic; Shot TR = 2800 ms; TI = 1098.6 ms; 256 shots, flip angle 8 degrees; TR = 8.3 ms; TE = 3.9 ms; SENSE acceleration in left-right direction 2; time-of-acquisition 5:35 minutes), while participants performed the calibration task.

### fMRI preprocessing

Results included in this manuscript come from preprocessing performed using fMRIPrep 20.2.3 (Esteban et al., 2019) which is based on Nipype 1.6.1 (Gorgolewski et al., 2011, 2018).

#### Anatomical data preprocessing

The T1-weighted (T1w) image was corrected for intensity non-uniformity (INU) with N4BiasFieldCorrection (Tustison et al., 2010), distributed with ANTs 2.3.3 (Avants et al., 2009), and used as T1w-reference throughout the workflow. The T1w-reference was then skull-stripped with a Nipype implementation of the antsBrainExtraction.sh workflow (from ANTs), using OASIS30ANTs as target template. Brain tissue segmentation of cerebrospinal fluid (CSF), white-matter (WM) and gray-matter (GM) was performed on the brain-extracted T1w using fast (FSL 5.0.9; Zhang et al., 2001). Brain surfaces were reconstructed using recon-all (FreeSurfer 6.0.1; Dale et al., 1999), and the brain mask estimated previously was refined with a custom variation of the method to reconcile ANTs-derived and FreeSurfer-derived segmentations of the cortical gray-matter of Mindboggle (Klein et al., 2017). Volume-based spatial normalization to one standard space (MNI152NLin2009cAsym) was performed through nonlinear registration with antsRegistration (ANTs 2.3.3), using brain-extracted versions of both T1w reference and the T1w template. The following template was selected for spatial normalization: ICBM 152 Nonlinear Asymmetrical template version 2009c (Fonov et al., 2009).

##### Functional data preprocessing

For each of the 6-18 BOLD runs found per participant (across all tasks and sessions), the following preprocessing was performed. First, a reference volume and its skull-stripped version were generated using a custom methodology of fMRIPrep. A B0-nonuniformity map (or fieldmap) was estimated based on two (or more) echo-planar imaging (EPI) references with opposing phase-encoding directions, with 3dQwarp Cox and Hyde (1997) (AFNI 20160207). Based on the estimated susceptibility distortion, a corrected EPI (echo-planar imaging) reference was calculated for a more accurate co-registration with the anatomical reference. The BOLD reference was then co-registered to the T1w reference using bbregister (FreeSurfer) which implements boundary-based registration (Greve & Fischl, 2009). Co-registration was configured with six degrees of freedom. Head-motion parameters with respect to the BOLD reference (transformation matrices, and six corresponding rotation and translation parameters) are estimated before any spatiotemporal filtering using mcflirt (FSL 5.0.9; Jenkinson et al., 2002). BOLD runs were slice-time corrected using 3dTshift from AFNI 20160207 (Cox & Hyde, 1997). The BOLD time-series were resampled onto the following surfaces (FreeSurfer reconstruction nomenclature): fsaverage, fsnative. The BOLD time-series (including slice-timing correction when applied) were resampled onto their original, native space by applying a single, composite transform to correct for head-motion and susceptibility distortions. These resampled BOLD time-series will be referred to as preprocessed BOLD in original space, or just preprocessed BOLD. The BOLD time-series were resampled into standard space, generating a preprocessed BOLD run in MNI152NLin2009cAsym space. First, a reference volume and its skull-stripped version were generated using a custom methodology of fMRIPrep. All resamplings can be performed with a single interpolation step by composing all the pertinent transformations (i.e. head-motion transform matrices, susceptibility distortion correction when available, and co-registrations to anatomical and output spaces). Gridded (volumetric) resamplings were performed using antsApplyTransforms (ANTs), configured with Lanczos interpolation to minimize the smoothing effects of other kernels (Lanczos, 1964). Non-gridded (surface) resamplings were performed using mri_vol2surf (FreeSurfer).

Many internal operations of fMRIPrep use Nilearn 0.6.2 (Abraham et al., 2014), mostly within the functional processing workflow. For more details of the pipeline, see the section corresponding to workflows in fMRIPrep’s documentation.

### fMRI analyses

To better understand the impact of cTBS on the neural representation of numbers, we used an encoding/decoding-modeling approach (Barretto-García et al., 2023; de Hollander, Grueschow, et al., 2024; van Bergen et al., 2015; Walker et al., 2020). We fitted a standard numerical receptive field (nPRF) model (de Hollander, Grueschow, et al., 2024; Harvey et al., 2013, 2020) that describes how a voxel 𝑖 non-linearly responds to specific numerosities 𝑠 𝑓(𝑠)→𝑥_𝑖_. Then, using a Bayesian framework, we also extended this encoding model with a multivariate likelihood function 𝑝(𝑋|𝑠) using a multivariate t-distribution: [𝑥_1_,.., 𝑥_𝑛_] ∼ 𝑓_1..𝑛_(𝑠) + ɛ with ɛ ∼ 𝑡(0, Σ, 𝑑), where Σ is the residual covariance and 𝑑 is the degrees-of-freedom of the t-distribution. Using an explicit likelihood function allows us to decode from trial-to-trial what was the presented payoff magnitude, as well as the acuity of the neural response.

The main fMRI analysis can roughly be split up in the following steps: 1) Fit a single-trial GLM to estimate trialwise measures of the response amplitude across voxels 2) Fit a numerical receptive field model (Harvey et al., 2013) to the response, 3) Fit a multivariate noise model to the residuals of the nPRF model in a leave-one-run-out cross-validation scheme (van Bergen et al., 2015), 4) Obtain a posterior estimate of the payoffs magnitudes of unseen data using the noise model and an inverted nPRF model.

#### Single trial estimates

We used the GLMSingle Python package (Prince et al., 2022) to obtain single-trial BOLD estimates. Briefly, the GLMSingle package uses cross-validation to do model selection over GLMs with a) a library of different hemodynamic response functions b) different L2-regularisation parameters to shrink the single trial estimates, combating the issue of correlated single trial regressors (Mumford et al., 2012). c) GLMSingle also obtains GLMDenoise (Kay et al., 2013) regressors based on the first *n* PCA components in a set of noise voxels. Noise voxels are defined as having low explained variance in the task-based GLM. The number of PCA components is selected via cross-validation.

As input to GLMSingle, we modeled the first and second payoff presentations separately. For the second payoff presentations, we modeled all trials with the same numerosity as being in the same condition, to aid GLMSingle with cross-validation (similar numerosities should have similar responses). After extensive analysis piloting, we chose to not include any additional confound regressors (e.g. motion parameters, RETROICOR parameters or aCompCorr regressors) in addition to the GLMDenoise confound regressors, as additional regressors did not lead to robustly increased (or even decreased) decoding accuracy and GLMDenoise. This is in line with earlier work on GLMDenoise (Kay et al., 2013) and the related aCompCorr approach (Behzadi et al., 2007).

#### Numerical receptive field modelling

We fitted a numerical population receptive field (nPRF) model to all voxels in the brain using the *braincoder* toolbox (de Hollander et al., 2020). We fitted a nPRF model for each session separately (so using 120 single trial estimates) for all sessions separately. The method is described in detail elsewhere (Barretto-García et al., 2023; Harvey et al., 2013), so we only briefly describe it here. First, we estimate the mean µ, standard deviation σ, amplitude 𝐴 and baseline 𝐵 of a log-normal receptive field for every voxel in the brain separately, to predict the BOLD response of these voxels to different numerosities.

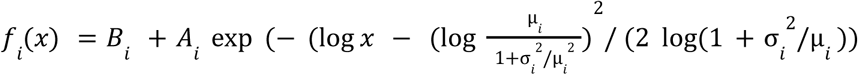

The second part of this equation is a parameterisation of the log-normal probability density function where µ and σ are the mean and standard deviation of the distribution in natural space. Although this parameterisation is somewhat exotic, it is highly useful when plotting the estimated preferred numerosities and their dispersion, for example on the cortical surface.

We first fit the model by using a grid-search: We correlate the single trial estimates for the first payoff stimulus presentation with the predictions of a large grid of 60 µ-s between 5 and 80 and 60 σ-s between 5 and 40. We then estimate 𝐼 and 𝐴 linear least-squares on the best-correlating µ and σ-parameters. Finally, we used gradient descent (Kingma & Ba, 2014) to refine parameters further. We then did the same procedure with a leave-one-run-out cross-validation scheme (so fit the model 6 times, always leaving one run out), to estimate the *cross-validate explained variance*, cvR2. Cross-validated R2 was used to distinguish ‘noise’ from ‘signal’ voxels. Specifically, within a 2cm-radius ROI around the targeted cluster, we selected all voxels that showed a cross-validated R2 larger than 0 for further inspection (i.e., parameter comparisons across conditions as well as decoding).

#### Decoding

After voxel selection, we used a leave-one-run-out cross validation scheme where the nPRF model was fitted to all runs but the test run, after which also a multivariate noise model was fitted to the residual signals. Specifically, we fitted the following covariance matrix:

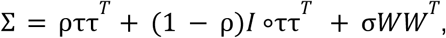

as well as the degrees-of-freedom of a 0-centred, multivariate t-distribution. Briefly, τ is a vector as its length the number of voxels (𝑛) in the ROI. It pertains to the standard deviation of the residuals of each voxel. Thus, ρ determines to which extent all voxels correlate with each other (ττ^𝑇^ is the covariance matrix of perfectly correlated voxels, whereas 𝐼 ∘ττ^𝑇^ corresponds to a perfectly diagonal matrix/spherical covariance). 𝑊 is a square matrix of 𝑛 ×𝑛. Each element 𝑊_𝑖,𝑗_ is the product of the receptive fields of voxels 𝑖 and 𝑗 across stimulus space (𝑓_𝑖_(𝑆) 𝑓_𝑗_(𝑆)^𝑇^ with 𝑆 being the entire stimulus space [7, 6,.., 112]. Thus, the free scalar parameter σ determines to which extent voxels with overlapping receptive fields have more correlated noise.

Once the noise model is fitted, we can now determine a likelihood function for any multivariate BOLD pattern 𝑋 and for any numerical stimulus 𝑠:

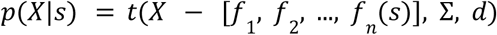

We assume a flat prior on all integers between 7 and 112, and can therefore evaluate this likelihood on all these integers and normalize the resulting probability mass function (pmf) to integrate to 1. We take the expected value of this pmf to be our estimate of the presented stimulus. We then correlated the actual stimulus numerosity with the estimated numerosity within every run and took the average correlation across all six runs to get the decoding performance for a given subject/session (Etzel et al., 2013).

## Code availability

All data of this study will be published on openneuro as soon as the study is published.

All the code used in this study can be found online on github: https://github.com/Gilles86/tms_risk/

Our computational cognitive models are implemented in the open-source Python library *bauer* https://github.com/ruffgroup/bauer/tree/main

The used nPRF model and decoding software algorithms are implemented in the open-source Python library *braincoder* https://braincoder-devs.github.io/

## Acknowledgements

We are grateful to C. Schnyder and K. Treiber at the Zurich Center for Neuroeconomics for their excellent assistance in recruitment and scanning. C.C.R. received funding from the University Research Priority Program ‘Adaptive Brain Circuits in Development and Learning’ at the University of Zurich and the Swiss National Science Foundation SNSF (grant no. 100019L-173248). G.d.H. was funded by the Dutch Research Council NWO (Rubicon grant no. 019.183SG.017/8O3B) and the University of Zurich (Forschungskredit grant no. K-33153-02-01)

## Footnotes

1 Risk-neutral probability (RNP) is one way to define the indifference point of a decision-maker and is the payout-probability that the risky option would have to have for a risk-neutral decision-maker to make the same choices as those empirically observed. Thus, a RNP below 55% corresponds to risk aversion, a RNP of 55% corresponds to risk-neutrality, and a RNP above 55% corresponds to risk-seeking (Barretto-García et al., 2023; de Hollander, Grueschow, et al., 2024; Khaw et al., 2020).

## References

Abraham, A., Pedregosa, F., Eickenberg, M., Gervais, P., Mueller, A., Kossaifi, J., Gramfort, A., Thirion, B., & Varoquaux, G. (2014). Machine learning for neuroimaging with scikit-learn. Frontiers in Neuroinformatics, 8, 14. 10.3389/fninf.2014.00014

Alserda, G. A. G., Dellaert, B. G. C., Swinkels, L., & Lecq, F. S. G. van der. (2019). Individual pension risk preference elicitation and collective asset allocation with heterogeneity. Journal of Banking & Finance, 101, 206–225. 10.1016/j.jbankfin.2019.02.014

Avants, B. B., Tustison, N., & Song, G. (2009). Advanced normalization tools (ANTS). 2.

Barretto-García, M., de Hollander, G., Grueschow, M., Polanía, R., Woodford, M., & Ruff, C. C. (2023). Individual risk attitudes arise from noise in neurocognitive magnitude representations. Nature Human Behaviour, 7(9), 1551–1567. 10.1038/s41562-023-01643-4

Bartra, O., McGuire, J. T., & Kable, J. W. (2013). The valuation system: A coordinate-based meta-analysis of BOLD fMRI experiments examining neural correlates of subjective value. NeuroImage, 76, 412–427. 10.1016/j.neuroimage.2013.02.063

Behzadi, Y., Restom, K., Liau, J., & Liu, T. T. (2007). A component based noise correction method (CompCor) for BOLD and perfusion based fMRI. NeuroImage, 37(1), 90–101. 10.1016/j.neuroimage.2007.04.042

Brown, R. I. F. (1986). Arousal and Sensation-Seeking Components in the General Explanation of Gambling and Gambling Addictions. International Journal of the Addictions, 21(9–10), 1001–1016. 10.3109/10826088609077251

Bueti, D., & Walsh, V. (2009). The parietal cortex and the representation of time, space, number and other magnitudes. Philosophical Transactions of the Royal Society B: Biological Sciences, 364(1525), 1831–1840. 10.1098/rstb.2009.0028

Busemeyer, J., & Townsend, J. (1993). Decision field theory: a dynamic-cognitive approach to decision making in an uncertain environment. 100(3).

Cai, Y., Hofstetter, S., Dijk, J. van, Zuiderbaan, W., Zwaag, W. van der, Harvey, B. M., & Dumoulin, S. O. (2021). Topographic numerosity maps cover subitizing and estimation ranges. Nature Communications, 12(1), 3374. 10.1038/s41467-021-23785-7

Cai, Y., Hofstetter, S., & Dumoulin, S. O. (2023). Nonsymbolic Numerosity Maps at the Occipitotemporal Cortex Respond to Symbolic Numbers. The Journal of Neuroscience, 43(16), 2950–2959. 10.1523/jneurosci.0687-22.2023

Capretto, T., Piho, C., Kumar, R., Westfall, J., Yarkoni, T., & Martin, O. A. (2022). Bambi: A Simple Interface for Fitting Bayesian Linear Models in Python. Journal of Statistical Software, 103(15), 1–29. 10.18637/jss.v103.i15

Chew, B., Hauser, T. U., Papoutsi, M., Magerkurth, J., Dolan, R. J., & Rutledge, R. B. (2019). Endogenous fluctuations in the dopaminergic midbrain drive behavioral choice variability. Proceedings of the National Academy of Sciences, 116(37), 18732–18737. 10.1073/pnas.1900872116

Coutlee, C. G., Kiyonaga, A., Korb, F. M., Huettel, S. A., & Egner, T. (2016). Reduced Risk-Taking following Disruption of the Intraparietal Sulcus. Frontiers in Neuroscience, 10, 588. 10.3389/fnins.2016.00588

Cox, R. W., & Hyde, J. S. (1997). Software tools for analysis and visualization of fMRI data. NMR in Biomedicine, 10(4-5), 171–178. 10.1002/(sici)1099-1492(199706/08)10:4/5<171::aid-nbm453>3.0.co;2-l

Dale, A. M., Fischl, B., & Sereno, M. I. (1999). Cortical Surface-Based Analysis I. Segmentation and Surface Reconstruction. NeuroImage, 9(2), 179–194. 10.1006/nimg.1998.0395

Dantas, A. M., Sack, A. T., Bruggen, E., Jiao, P., & Schuhmann, T. (2021). Reduced risk-taking behavior during frontal oscillatory theta band neurostimulation. Brain Research, 1759, 147365. 10.1016/j.brainres.2021.147365

Dantas, A. M., Sack, A. T., Bruggen, E., Jiao, P., & Schuhmann, T. (2023). Modulating risk-taking behavior with theta-band tACS. NeuroImage, 120422. 10.1016/j.neuroimage.2023.120422

Dehaene, S. (2011). The number sense: How the mind creates mathematics. Oxford University Press.

Dehaene, S., & Cohen, L. (1991). Two mental calculation systems: A case study of severe acalculia with preserved approximation. Neuropsychologia, 29(11), 1045–1074. 10.1016/0028-3932(91)90076-k

Dehaene, S., Dehaene-Lambertz, G., & Cohen, L. (1998). Abstract representations of numbers in the animal and human brain. Trends in Neurosciences, 21(8), 355–361. 10.1016/s0166-2236(98)01263-6

Dehaene, S., Piazza, M., Pinel, P., & Cohen, L. (2003). Three Parietal Circuits for Number Processing. Cognitive Neuropsychology, 20(3–6), 487–506. 10.1080/02643290244000239

de Hollander, G., Grueschow, M., Hennel, F., & Ruff, C. C. (2024). Rapid Changes in Risk Preferences Originate from Bayesian Inference on Parietal Magnitude Representations. BioRxiv, 2024.08.23.609296. 10.1101/2024.08.23.609296

de Hollander, G., Renkert, M. F., Knapen, T. H., & Ruff, C. C. (2020). Braincoder: Encoding models for fMRI implemented in tensorflow. https://braincoder-devs.github.io/

de Hollander, G., Renkert, M. F., & Ruff, C. C. (2024). Bauer: Bayesian Estimation of Perceptual, Numerical and Risky Judgements. https://github.com/ruffgroup/bauer/tree/main

Drichoutis, A. C., & Lusk, J. L. (2016). What can multiple price lists really tell us about risk preferences? Journal of Risk and Uncertainty, 53(2–3), 89–106. 10.1007/s11166-016-9248-5

Dumoulin, S. O., Fracasso, A., Zwaag, W. van der, Siero, J. C. W., & Petridou, N. (2018). Ultra-high field MRI: Advancing systems neuroscience towards mesoscopic human brain function. NeuroImage, 168(Nat. Neurosci. 6 2003), 345–357. 10.1016/j.neuroimage.2017.01.028

Eger, E., Michel, V., Thirion, B., Amadon, A., Dehaene, S., & Kleinschmidt, A. (2009). Deciphering Cortical Number Coding from Human Brain Activity Patterns. Current Biology, 19(19), 1608–1615. 10.1016/j.cub.2009.08.047

Eger, E., Sterzer, P., Russ, M. O., Giraud, A.-L., & Kleinschmidt, A. (2003). A Supramodal Number Representation in Human Intraparietal Cortex. Neuron, 37(4), 719–726. 10.1016/s0896-6273(03)00036-9

Esteban, O., Markiewicz, C. J., Blair, R. W., Moodie, C. A., Isik, A. I., Erramuzpe, A., Kent, J. D., Goncalves, M., DuPre, E., Snyder, M., Oya, H., Ghosh, S. S., Wright, J., Durnez, J., Poldrack, R. A., & Gorgolewski, K. J. (2019). fMRIPrep: a robust preprocessing pipeline for functional MRI. Nature Methods, 16(1), 111–116. 10.1038/s41592-018-0235-4

Etzel, J. A., Zacks, J. M., & Braver, T. S. (2013). Searchlight analysis: Promise, pitfalls, and potential. NeuroImage, 78, 261–269. 10.1016/j.neuroimage.2013.03.041

FeldmanHall, O., Glimcher, P., Baker, A. L., & Phelps, E. A. (2016). Emotion and Decision-Making Under Uncertainty: Physiological Arousal Predicts Increased Gambling During Ambiguity but Not Risk. Journal of Experimental Psychology: General, 145(10), 1255–1262. 10.1037/xge0000205

Fiedler, S., & Glöckner, A. (2012). The Dynamics of Decision Making in Risky Choice: An Eye-Tracking Analysis. Frontiers in Psychology, 3, 335. 10.3389/fpsyg.2012.00335

Fonov, V., Evans, A., McKinstry, R., Almli, C., & Collins, D. (2009). Unbiased nonlinear average age-appropriate brain templates from birth to adulthood. NeuroImage, 47, S102. 10.1016/s1053-8119(09)70884-5

Foucault, C., Bounmy, T., Demortain, S., Thiririon, B., Eger, E., & Meyniel, F. (2024). A nonlinear code for event probability in the human brain. BioRxiv, 2024.02.28.582455. 10.1101/2024.02.28.582455

Frydman, C., & Jin, L. J. (2021). Efficient Coding and Risky Choice. The Quarterly Journal of Economics, 137(1), 161–213. 10.1093/qje/qjab031

Gates, N. J. S. and M. W. and C. H. and broessli and P. Q. and S. S. and A. P. and B. B. and C. D.-P. and H. K. and K. L. and M. H. and M. H.-D. and M. K. and T. (2023). pydata/patsy: v0.5.4. 10.5281/zenodo.10233918

Gelman, A., Carlin, J. B., Stern, H. S., Dunson, D. B., Vehtari, A., & Rubin, D. B. (2013). Bayesian data analysis. Chapman and Hall/CRC.

Gianotti, L. R. R., Knoch, D., Faber, P. L., Lehmann, D., Pascual-Marqui, R. D., Diezi, C., Schoch, C., Eisenegger, C., & Fehr, E. (2008). Tonic Activity Level in the Right Prefrontal Cortex Predicts Individuals’ Risk Taking. Psychological Science, 20(1), 33–38. 10.1111/j.1467-9280.2008.02260.x

Gilaie-Dotan, S., Tymula, A., Cooper, N., Kable, J. W., Glimcher, P. W., & Levy, I. (2014). Neuroanatomy Predicts Individual Risk Attitudes. The Journal of Neuroscience, 34(37), 12394–12401. 10.1523/jneurosci.1600-14.2014

Gorgolewski, K. J., Burns, C. D., Madison, C., Clark, D., Halchenko, Y. O., Waskom, M. L., & Ghosh, S. S. (2011). Nipype: A Flexible, Lightweight and Extensible Neuroimaging Data Processing Framework in Python. Frontiers in Neuroinformatics, 5, 13. 10.3389/fninf.2011.00013

Gorgolewski, K. J., Esteban, O., Markiewicz, C. J., Ziegler, E., Ellis, D. G., Notter, M. P., Jarecka, D., Johnson, H., Burns, C., Manhães-Savio, A., Hamalainen, C., Yvernault, B., Salo, T., Jordan, K., Goncalves, M., Waskom, M., Clark, D., Wong, J., Loney, F., … Perkins, L. (2018). Nipype. Software. Zenodo. 10.5281/Zenodo, 596855.

Green, D. M., & Swets, J. A. (1966). Signal detection theory and psychophysics (Vol. 1). Wiley New York.

Greve, D. N., & Fischl, B. (2009). Accurate and robust brain image alignment using boundary-based registration. 48(1). 10.1016/j.neuroimage.2009.06.060

Griffiths, M. D. (1990). The cognitive psychology of gambling. Journal of Gambling Studies, 6(1), 31–42. 10.1007/bf01015747

Harvey, B. M. (2016). Quantity Cognition: Numbers, Numerosity, Zero and Mathematics. Current Biology, 26(10), R419–R421. 10.1016/j.cub.2016.03.059

Harvey, B. M., Dumoulin, S. O., Fracasso, A., & Paul, J. M. (2020). A Network of Topographic Maps in Human Association Cortex Hierarchically Transforms Visual Timing-Selective Responses. Current Biology, 30(8), 1424–1434.e6. 10.1016/j.cub.2020.01.090

Harvey, B. M., Fracasso, A., Petridou, N., & Dumoulin, S. O. (2015). Topographic representations of object size and relationships with numerosity reveal generalized quantity processing in human parietal cortex. Proceedings of the National Academy of Sciences, 112(44), 13525–13530. 10.1073/pnas.1515414112

Harvey, B. M., Klein, B. P., Petridou, N., & Dumoulin, S. O. (2013). Topographic Representation of Numerosity in the Human Parietal Cortex. Science, 341(6150), 1123–1126. 10.1126/science.1239052

Heerema, R., Carrillo, P., Daunizeau, J., Vinckier, F., & Pessiglione, M. (2023). Mood fluctuations shift cost–benefit tradeoffs in economic decisions. Scientific Reports, 13(1), 18173. 10.1038/s41598-023-45217-w

Hoffman, M. D., & Gelman, A. (2011). The No-U-Turn Sampler: Adaptively Setting Path Lengths in Hamiltonian Monte Carlo. Journal of Machine Learning Research, 15, 1593–1623.

Hollander, G. de, Grueschow, M., Hennel, F., & Ruff, C. C. (2024). Rapid changes in risk attitudes originate from Bayesian inference on noisy neural magnitude representations.

Huang, Y.-Z., Edwards, M. J., Rounis, E., Bhatia, K. P., & Rothwell, J. C. (2005). Theta Burst Stimulation of the Human Motor Cortex. Neuron, 45(2), 201–206. 10.1016/j.neuron.2004.12.033

Jahedi, S., Deck, C., & Ariely, D. (2017). Arousal and economic decision making. Journal of Economic Behavior & Organization, 134, 165–189. 10.1016/j.jebo.2016.10.008

Jenkinson, M., Bannister, P., Brady, M., & Smith, S. (2002). Improved optimization for the robust and accurate linear registration and motion correction of brain images. Neuroimage, 17(2), 825–841.

Kay, K. N., Rokem, A., Winawer, J., Dougherty, R. F., & Wandell, B. A. (2013). GLMdenoise: a fast, automated technique for denoising task-based fMRI data. Frontiers in Neuroscience, 7, 247. 10.3389/fnins.2013.00247

Khaw, M. W., Li, Z., & Woodford, M. (2020). Cognitive Imprecision and Small-Stakes Risk Aversion. The Review of Economic Studies, 88(4), 1979–2013. 10.1093/restud/rdaa044

Kingma, D. P., & Ba, J. (2014). Adam: A Method for Stochastic Optimization. ArXiv. 10.48550/arxiv.1412.6980

Klein, A., Ghosh, S. S., Bao, F. S., Giard, J., Häme, Y., Stavsky, E., Lee, N., Rossa, B., Reuter, M., Neto, E. C., & Keshavan, A. (2017). Mindboggling morphometry of human brains. PLoS Computational Biology, 13(2), e1005350. 10.1371/journal.pcbi.1005350

Lanczos, C. (1964). Evaluation of Noisy Data. Journal of the Society for Industrial and Applied Mathematics Series B Numerical Analysis, 1(1), 76–85. 10.1137/0701007

Landau, A. N., Schreyer, H. M., van Pelt, S., & Fries, P. (2015). Distributed Attention Is Implemented through Theta-Rhythmic Gamma Modulation. Current Biology, 25(17), 2332–2337. 10.1016/j.cub.2015.07.048

Lasne, G., Piazza, M., Dehaene, S., Kleinschmidt, A., & Eger, E. (2019). Discriminability of numerosity-evoked fMRI activity patterns in human intra-parietal cortex reflects behavioral numerical acuity. Cortex, 114, 90–101. 10.1016/j.cortex.2018.03.008

Lee, M. D., & Wagenmakers, E.-J. (2014). Bayesian Cognitive Modeling: A Practical Course. Cambridge university press. 10.1017/cbo9781139087759

Levy, D. J., & Glimcher, P. W. (2012). The root of all value: a neural common currency for choice. Current Opinion in Neurobiology, 22(6), 1027–1038. 10.1016/j.conb.2012.06.001

Mata, R., Frey, R., Richter, D., Schupp, J., & Hertwig, R. (2018). Risk Preference: A View from Psychology. Journal of Economic Perspectives, 32(2), 155–172. 10.1257/jep.32.2.155

Miedl, S. F., Peters, J., & Büchel, C. (2012). Altered Neural Reward Representations in Pathological Gamblers Revealed by Delay and Probability Discounting. Archives of General Psychiatry, 69(2), 177–186. 10.1001/archgenpsychiatry.2011.1552

Mohr, P. N. C., Biele, G., & Heekeren, H. R. (2010). Neural Processing of Risk. The Journal of Neuroscience, 30(19), 6613–6619. 10.1523/jneurosci.0003-10.2010

Mumford, J. A., Turner, B. O., Ashby, F. G., & Poldrack, R. A. (2012). Deconvolving BOLD activation in event-related designs for multivoxel pattern classification analyses. NeuroImage, 59(3), 2636–2643. 10.1016/j.neuroimage.2011.08.076

Olschewski, S., & Rieskamp, J. (2021). Distinguishing three effects of time pressure on risk taking: Choice consistency, risk preference, and strategy selection. Journal of Behavioral Decision Making, 34(4), 541–554. 10.1002/bdm.2228

Olschewski, S., Rieskamp, J., & Scheibehenne, B. (2018). Taxing Cognitive Capacities Reduces Choice Consistency Rather Than Preference: A Model-Based Test. Journal of Experimental Psychology: General, 147(4), 462–484. 10.1037/xge0000403

Olschewski, S., Sirotkin, P., & Rieskamp, J. (2022). Empirical underidentification in estimating random utility models: The role of choice sets and standardizations. British Journal of Mathematical and Statistical Psychology, 75(2), 252–292. 10.1111/bmsp.12256

Paliwal, S., Mosley, P. E., Breakspear, M., Coyne, T., Silburn, P., Aponte, E., Mathys, C., & Stephan, K. E. (2019). Subjective estimates of uncertainty during gambling and impulsivity after subthalamic deep brain stimulation for Parkinson’s disease. Scientific Reports, 9(1), 14795. 10.1038/s41598-019-51164-2

Paliwal, S., Petzschner, F. H., Schmitz, A. K., Tittgemeyer, M., & Stephan, K. E. (2014). A model-based analysis of impulsivity using a slot-machine gambling paradigm. Frontiers in Human Neuroscience, 8, 428. 10.3389/fnhum.2014.00428

Panidi, K., Vorobiova, A. N., Feurra, M., & Klucharev, V. (2023). Posterior parietal cortex is causally involved in reward valuation but not in probability weighting during risky choice. Cerebral Cortex, 34(1), bhad446. 10.1093/cercor/bhad446

Patil, A., Huard, D., & Fonnesbeck, C. J. (2010). PyMC: Bayesian Stochastic Modelling in Python. Journal of Statistical Software, 35(4), 1–81.

Pedroni, A., Frey, R., Bruhin, A., Dutilh, G., Hertwig, R., & Rieskamp, J. (2017). The risk elicitation puzzle. Nature Human Behaviour, 1(11), 803–809. 10.1038/s41562-017-0219-x

Peters, E., Västfjäll, D., Slovic, P., Mertz, C. K., Mazzocco, K., & Dickert, S. (2005). Numeracy and Decision Making. Psychological Science, 17(5), 407–413. 10.1111/j.1467-9280.2006.01720.x

Petzschner, F. H., Glasauer, S., & Stephan, K. E. (2015). A Bayesian perspective on magnitude estimation. Trends in Cognitive Sciences, 19(5), 285–293. 10.1016/j.tics.2015.03.002

Pfeffer, T., Keitel, C., Kluger, D. S., Keitel, A., Russmann, A., Thut, G., Donner, T. H., & Gross, J. (2022). Coupling of pupil- and neuronal population dynamics reveals diverse influences of arousal on cortical processing. ELife, 11, e71890. 10.7554/elife.71890

Poldrack, R. A. (2017). Precision Neuroscience: Dense Sampling of Individual Brains. 95(4). 10.1016/j.neuron.2017.08.002

Pouget, A., Beck, J. M., Ma, W. J., & Latham, P. E. (2013). Probabilistic brains: knowns and unknowns. Nature Neuroscience, 16(9), 1170–1178. 10.1038/nn.3495

Prat-Carrabin, A., & Gershman, S. J. (2024). Bayes vs. Weber: how to break a law of psychophysics. BioRxiv, 2024.08.08.607196. 10.1101/2024.08.08.607196

Prat-Carrabin, A., & Woodford, M. (2022). Efficient coding of numbers explains decision bias and noise. Nature Human Behaviour, 6(8), 1142–1152. 10.1038/s41562-022-01352-4

Prince, J. S., Charest, I., Kurzawski, J. W., Pyles, J. A., Tarr, M. J., & Kay, K. N. (2022). Improving the accuracy of single-trial fMRI response estimates using GLMsingle. ELife, 11, e77599. 10.7554/elife.77599

Rogers, P. (1998). The Cognitive Psychology of Lottery Gambling: A Theoretical Review. Journal of Gambling Studies, 14(2), 111–134. 10.1023/a:1023042708217

Sack, A. T., Kadosh, R. C., Schuhmann, T., Moerel, M., Walsh, V., & Goebel, R. (2009). Optimizing Functional Accuracy of TMS in Cognitive Studies: A Comparison of Methods. Journal of Cognitive Neuroscience, 21(2), 207–221. 10.1162/jocn.2009.21126

Sacré, P., Kerr, M. S. D., Subramanian, S., Fitzgerald, Z., Kahn, K., Johnson, M. A., Niebur, E., Eden, U. T., González-Martínez, J. A., Gale, J. T., & Sarma, S. V. (2019). Risk-taking bias in human decision-making is encoded via a right–left brain push–pull system. Proceedings of the National Academy of Sciences, 116(4), 201811259. 10.1073/pnas.1811259115

Sharp, C., Monterosso, J., & Montague, P. R. (2012). Neuroeconomics: A Bridge for Translational Research. Biological Psychiatry, 72(2), 87–92. 10.1016/j.biopsych.2012.02.029

Spurrier, M., & Blaszczynski, A. (2014). Risk Perception in Gambling: A Systematic Review. Journal of Gambling Studies, 30(2), 253–276. 10.1007/s10899-013-9371-z

Spyropoulos, G., Bosman, C. A., & Fries, P. (2018). A theta rhythm in macaque visual cortex and its attentional modulation. Proceedings of the National Academy of Sciences, 115(24), E5614–E5623. 10.1073/pnas.1719433115

Stanton, S. J., Reeck, C., Huettel, S. A., & LaBar, K. S. (2014). Effects of induced moods on economic choices. Judgment and Decision Making, 9(2), 167–175. 10.1017/s1930297500005532

Stewart, N., Chater, N., & Brown, G. D. A. (2006). Decision by sampling. Cognitive Psychology, 53(1), 1–26. 10.1016/j.cogpsych.2005.10.003

Tustison, N. J., Avants, B. B., Cook, P. A., Zheng, Y., Egan, A., Yushkevich, P. A., & Gee, J. C. (2010). N4ITK: Improved N3 Bias Correction. IEEE Transactions on Medical Imaging, 29(6), 1310–1320. 10.1109/tmi.2010.2046908

van Bergen, R. S., Ma, W. J., Pratte, M. S., & Jehee, J. F. M. (2015). Sensory uncertainty decoded from visual cortex predicts behavior. Nature Neuroscience, 18(12), 1728. 10.1038/nn.4150

Vehtari, A., Gelman, A., & Gabry, J. (2017). Practical Bayesian model evaluation using leave-one-out cross-validation and WAIC. Statistics and Computing, 27(5), 1413–1432. 10.1007/s11222-016-9696-4

Walasek, L., & Stewart, N. (2015). How to Make Loss Aversion Disappear and Reverse: Tests of the Decision by Sampling Origin of Loss Aversion. Journal of Experimental Psychology: General, 144(1), 7–11. 10.1037/xge0000039

Walker, E. Y., Cotton, R. J., Ma, W. J., & Tolias, A. S. (2020). A neural basis of probabilistic computation in visual cortex. Nature Neuroscience, 23(1), 122–129. 10.1038/s41593-019-0554-5

Walsh, V. (2003). A theory of magnitude: common cortical metrics of time, space and quantity. Trends in Cognitive Sciences, 7(11), 483–488. 10.1016/j.tics.2003.09.002

Wang, L., Mruczek, R. E. B., Arcaro, M. J., & Kastner, S. (2015). Probabilistic Maps of Visual Topography in Human Cortex. Cerebral Cortex, 25(10), 3911–3931. 10.1093/cercor/bhu277

Warrington, E. K., & James, M. (1967). Tachistoscopic number estimation in patients with unilateral cerebral lesions. Journal of Neurology, Neurosurgery & Psychiatry, 30(5), 468. 10.1136/jnnp.30.5.468

Whitmarsh, S., Gitton, C., Jousmäki, V., Sackur, J., & Tallon-Baudry, C. (2022). Neuronal correlates of the subjective experience of attention. European Journal of Neuroscience, 55(11–12), 3465–3482. 10.1111/ejn.15395

Woodford, M. (2020). Modeling Imprecision in Perception, Valuation, and Choice. Annual Review of Economics, 12(1), 1–23. 10.1146/annurev-economics-102819-040518

Zhang, Y., Brady, M., & Smith, S. (2001). Segmentation of Brain MR Images Through a Hidden Markov Random Field Model and the Expectation-Maximization Algorithm. IEEE Transactions on Medical Imaging, 20(1), 45. 10.1109/42.906424

