## Supplementary figure 1 for "Risk preferences causally rely on parietal magnitude representations: Evidence from combined TMS-fMRI"

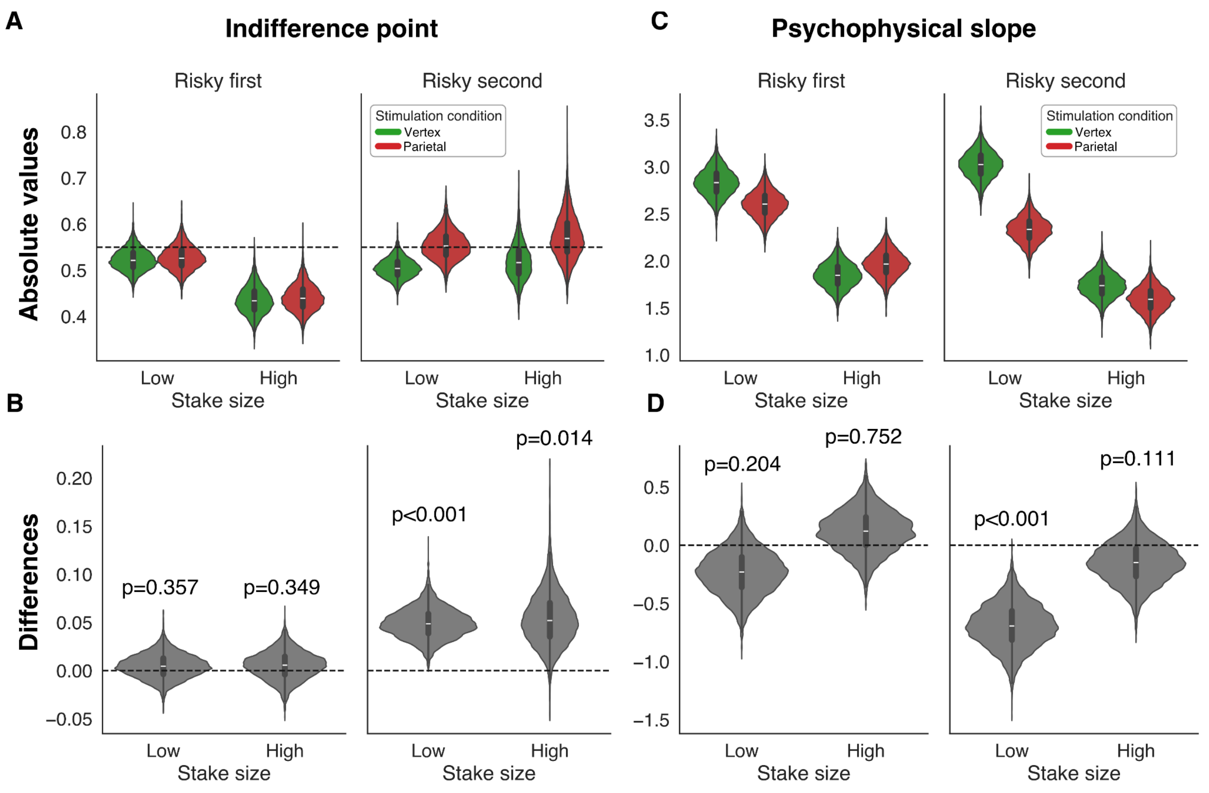


**Supplementary Figure 1: A)** Estimated average indifference points for low and high average stake sizes across different presentation orders. **B)** Differences in group average indifference points between TMS conditions. Notably, when the risky option is presented second, indifference points shift toward risk-seeking behavior, particularly for low stakes. **C)** Estimated average choice consistency, represented by the slope of the log risky-to-safe ratio on choice probability in a probit model. **D)** Differences in group average choice consistency estimates between TMS conditions. Choice consistency is significantly altered only when the risky option is presented second, with the effect being most pronounced for low stakes.
